# Universal functionalization of extracellular vesicles with nanobody adapters

**DOI:** 10.64898/2026.02.28.708726

**Authors:** Andrea Galisova, Jiri Zahradnik, Ester Merunkova, Dominik Havlicek, Josef Uskoba, Ziv Porat, Daniel Jirak

**Affiliations:** Institute for Clinical and Experimental Medicine, Videnska 1958/9, Prague, Czech Republic; The First Faculty of Medicine, Charles University, BIOCEV, Prumyslova 595, Vestec, Prague-region, Czech Republic; Institute of Biophysics, The First Faculty of Medicine, Charles University, Salmovska 1, Prague, Czech Republic; BioTech a.s., Kramolinska 955, Prague, Czech Republic; Life Sciences Core Facilities, Weizmann Institute of Science, Rehovot, Israel

**Keywords:** extracellular vesicles, EVs, cancer, drug delivery

## Abstract

Extracellular vesicles (EVs) have emerged as a powerful platform for targeted therapies due to their intrinsic capacity for intercellular communication and low immunogenicity. In addition to their desirable natural properties, EVs can be engineered to programmably display targeting moieties on their surface, leading to enhanced specificity. Current methods for EV engineering rely on genetic engineering of parental cells, which is robust but labor-intensive due to the requirement to generate stable cell lines for each targeting protein. To address this hurdle, we introduce the Nanobody-Tag-Ligand system (NaTaLi), in which anti-ALFA tag nanobodies are anchored to the EV surface, enabling flexible and nearly covalent attachment of ALFA-tagged proteins. Crucially, NaTaLi allows stable and uniform functionalization of isolated EVs with any tagged protein, removing the need for further mammalian cell engineering. We demonstrate that NaTaLi enables simultaneous display of multiple functional moieties, allowing for precise tunability. In a murine model of breast cancer, NaTaLi-engineered EVs exhibited specific, high-efficiency delivery to tumor cells *in vivo*. Thus, NaTaLi is a versatile, plug-and-play system that may accelerate the development of targeted EV therapeutics and open the door to readily engineering complex, multispecific EVs.

**Graphical Abstract:** A schematic representation of the NaTaLi delivery system. EVs are engineered to display ALFA nanobodies on their surface (ALFA-EVs). ALFA-tagged proteins of choice are isolated and purified from bacteria. Mixing ALFA-EVs with ALFA-tagged proteins creates EVs functionalized with proteins of choice. For examples, ALFA-EVs can be functionalized with tumor-targeting proteins for *in vivo* targeting of tumors.

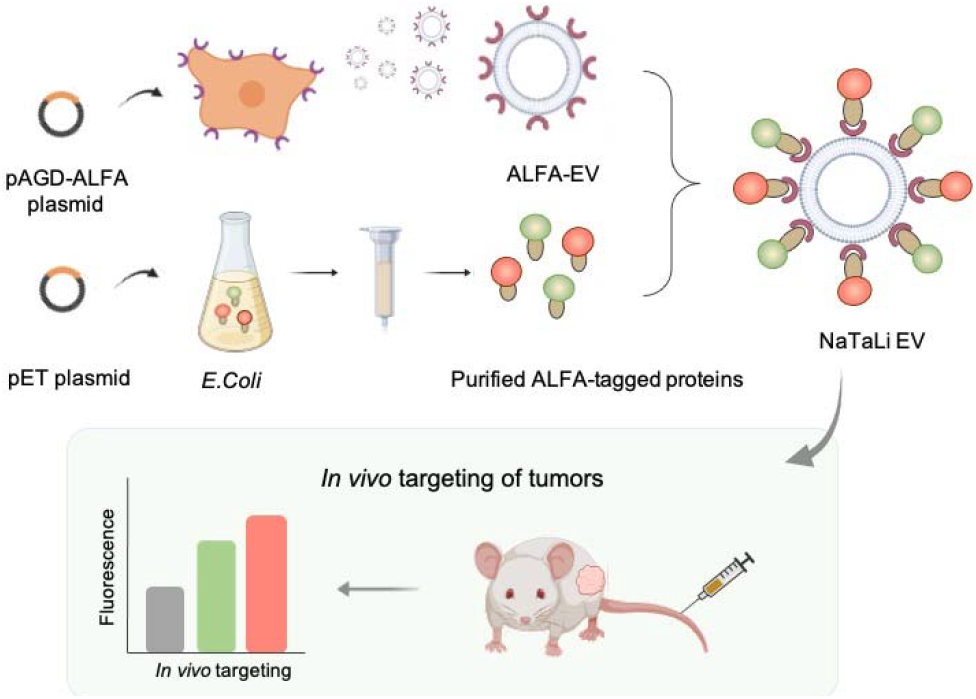

## 1. Introduction

Surface functionalization of extracellular vesicles (EVs) is an emerging tool for targeted drug delivery. EVs are particularly promising cell-free carries of therapeutic agents due to their natural role in cell-cell communication^1–3^, biocompatibility, and intrinsic ability to transport cargoes^4–6^. In addition, EVs can be modified to display or carry specialized proteins, enabling tissue-specific targeting^7,8^, and can be loaded with therapeutic agents, such as proteins, RNA or small drug molecules^6,9–12^. Importantly, EVs naturally cross multiple biological barriers, including the blood-brain-barrier^8^, allowing them to deliver cargo throughout the body.

One promising approach for the functionalization of EVs is to display specialized proteins on their surface. This can be achieved in principle by physical association of the protein of interest to the EV surface through a phospholipid anchor or chemical modification of the EV surface. However, molecules added to the EV surface after purification by chemical coupling^13^ or non-covalent addition into the phospholipid membrane can disrupt EV surface composition or dissociate from EVs *in vivo*^14,15^. An alternative strategy is to genetically fuse the protein of interest to a membrane-targeting motif and introduce the DNA-encoding for the fusion protein into producer cells. The producer cells then express the fusion protein, which is naturally inserted into the membrane during EV formation and displayed on the EV surface.

Current techniques for genetic engineering of EVs create one producer cell type per targeting protein. Although robust, these approaches are time-consuming and the amount of work scales with the number of proteins tested. Recent approaches reduce the amount of work necessary by displaying a docking moiety, such as an antibody Fc-binding domain^9,16^. This approach enables targeting of EVs to cancer cells through display of cancer-associated antibodies^9^, but is applicable only to antibodies or proteins with an Fc tag. Moreover, Fc-binding proteins have affinities in the low nM range, suggesting there is likely noticeable dissociation *in vivo*. Another emerging system utilizes fluorescein-mediated functionalization of EVs^17^ and represents a versatile platform, however its application has been demonstrated exclusively for antibodies and the expression levels were limited. We reasoned that these issues could be addressed by displaying a specialized nanobody with high expression levels and high affinity to a universal protein tag on the surface of EVs.

We engineered a stable parental cell line that produces EVs displaying the engineered ALFA nanobody on their surface^18^. ALFA nanobody is a low picomolar affinity nanobody that specifically recognizes the monomeric short protein ALFA tag^19^ that can easily be fused to any protein^20,21^. ALFA nanobody binding to the ALFA tag is very strong under native conditions^19^, enabling near covalent binding and therefore long-term stability. In addition to its binding properties, the ALFA nanobody was engineered to ensure efficient intracellular trafficking and robust surface expression^18^, making it an ideal target for protein display. Finally, ALFA nanobody expression and ALFA tag binding are constant, meaning that display of different proteins should also be constant. Thus, ALFA-EVs can be stably functionalized with any protein fused to an ALFA tag easily produced in bacteria without the need for further mammalian cell engineering.

In this study, we developed and optimized the Nanobody-Tag-Ligand EV system (NaTaLi) and demonstrated it enables *in vivo* targeting of EVs toward various cancer biomarkers. Crucially, we show that NaTaLi enables display of multiple proteins simultaneously, paving the way towards engineering of multispecific EVs or EVs endowed with several desirable properties.

## 2. Results

### 2.1 Functional display of the ALFA nanobody on cell membranes

We first established a stable cell line exhibiting high ALFA nanobody surface expression (HEK293-ALFA) by transfecting HEK293 cells with an ALFA nanobody-expressing plasmid (**Fig. S1**) and selecting transfected cells using puromycin selection and fluorescence-activated cell sorting (FACS, **Fig. 1A, Fig. S2A, B**). Western blot analysis confirmed the expression of ALFA nanobodies on the HEK293-ALFA cells (**Fig. 1J, Fig. S3A**). We verified that HEK293-ALFA cells bind strongly to two different fluorescent ALFA-tagged proteins (TP), mNeonGreen (mNeonTP) and dTomato (dTomatoTP), by both fluorescent microscopy and flow cytometry (**Fig. 1B-E**). Both fluorescent proteins showed distinct localization on the cell surface (**Fig. 1B, E and insets**), suggesting binding to correctly displayed nanobodies. Fluorescent proteins fused with control tags did not bind to HEK293-ALFA (flow cytometry histograms in **Fig. 1C, F** and quantification in **Fig. 1D, G**) confirming HE293-ALFA cells bind only ALFA-tagged proteins. We additionally verified that HEK293-ALFA cells specifically bind cancer-targeting peptides fused to an ALFA tag and a fluorescent protein (**Fig. S4C, D**). Thus, we concluded that HEK293-ALFA cells specifically bind to ALFA-tagged proteins.

**Fig. 1.**
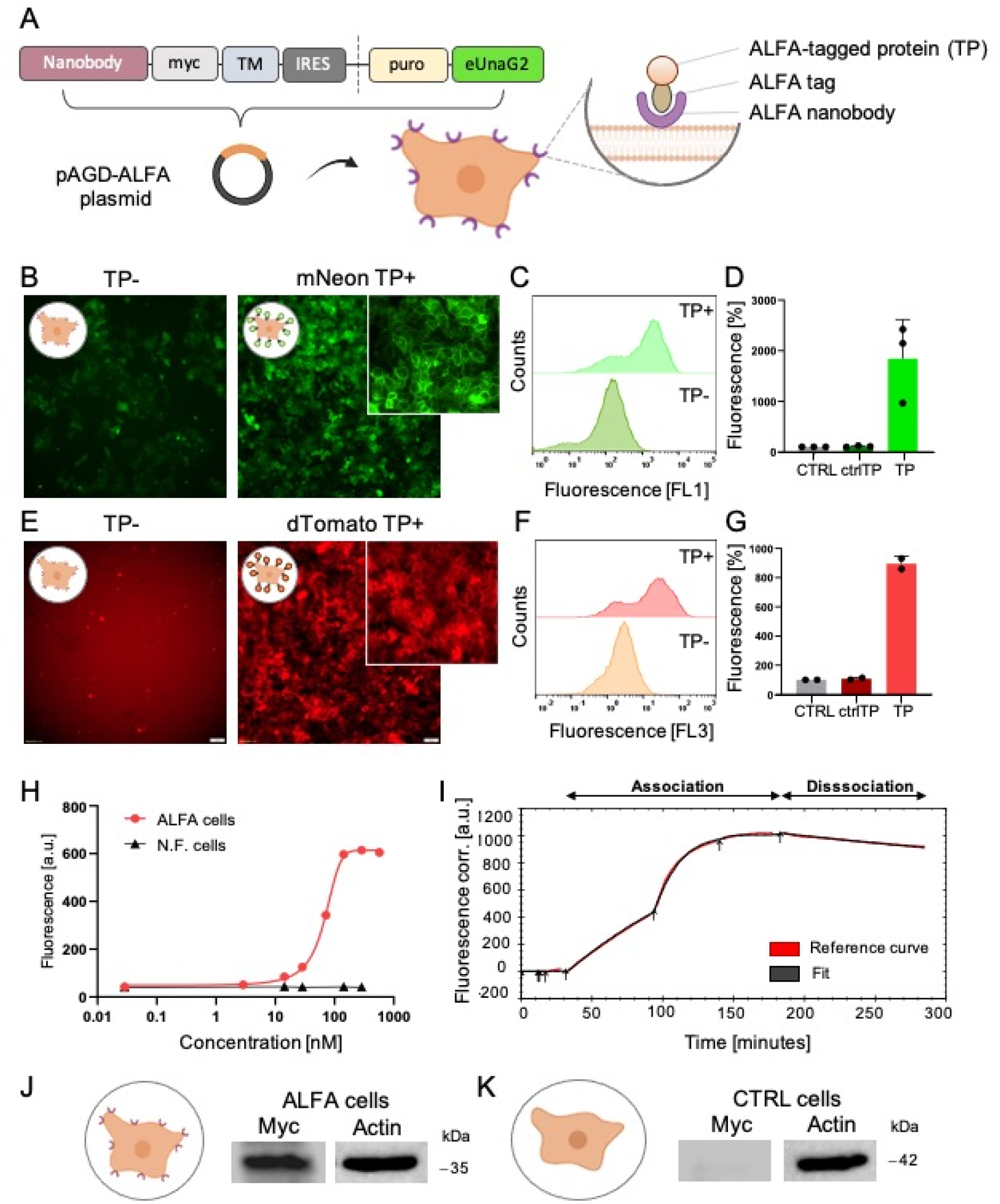
Characterization of ALFA nanobody-expressing cells. **(A)** HEK293 cells are transfected with an expression plasmid for the display of ALFA nanobodies on cell membrane surface. Microscopy images of ALFA nanobody cells with clicked ALFA-tagged proteins (TP) **(B)** mNeon and **(E)** dTomato on their surface. Flow cytometry histograms of ALFA nanobody cells with **(C)** mNeonTP and **(F)** dTomatoTP. Quantification of fluorescence signal at a single-cell level by flow cytometry **(D)** for mNeonTP and **(G)** for dTomatoTP. CtrlTP is a fluorescent protein fused with a control tag, TP is a fluorescent protein fused with the ALFA tag. The fluorescence signal was normalized to the signal of the control (CTRL) cells (without addition of proteins). **(H)** Binding of mNeonTP protein to ALFA nanobody cells measured by flow cytometry. **(I)** Association and dissociation of mNeonTP protein to ALFA cells measured by Ligand Tracer. Protein expression in **(J)** ALFA-expressing cells and **(K)** control non-functionalized (CTRL) HEK293 cells analyzed by western blot.

We next set out to measure the strength of HEK293-ALFA cells binding to ALFA-tagged proteins using several approaches. First, we incubated HEK293-ALFA cells with various concentrations of mNeonTP and measured binding using flow cytometry, revealing specific and concentration-dependent binding of tagged proteins to HEK293-ALFA cells (**Fig. 1H**, histograms shown in **Fig. S2C**). Next, we applied Ligand Tracer to measure the affinity of ALFA nanobodies expressed on the cell surface to ALFA-tagged proteins in real time. The nanobodies exhibited rapid association (k_a_ = 2.42×10^4^ M^-1^s^-1^) and very slow dissociation (k_d_ = 1.89×10^-5^ s^-1^), resulting in a *K*_*D*_ of 782 pM. These results confirm HEK293-ALFA cells bind strongly to ALFA-tagged proteins and suggest binding results in long-lasting functionalization (**Fig. 1I, Fig. S3**).

### 2.2 Flexible and robust functionalization of EVs with ALFA-tagged proteins

Having shown that HEK293-ALFA cells display ALFA-nanobodies on their surface, we next evaluated if EVs produced by HEK293-ALFA cells also display ALFA-nanobodies. EVs displaying ALFA nanobodies were isolated from HEK293-ALFA cells (ALFA-EVs) and validated using western blots, nanoparticle tracking analysis (NTA) and cryogenic electron microscopy (cryo-EM) according to the MISEV 2023 criteria^22^. Western blot analysis confirmed the presence of the ALFA nanobody on EVs, and the EV-specific marker CD81 was present in both ALFA-EVs and control EVs (**Fig. 2D, Fig. S5**). ALFA-EVs were then functionalized with ALFA-tagged proteins produced in *E*.*Coli* (**Fig. S4A**). Nanoflow cytometry verified at a single-vesicle level that ALFA-EVs bind to an ALFA-tagged fluorescent protein, while control EVs do not (**Fig. 2E, F**).

**Fig. 2.**
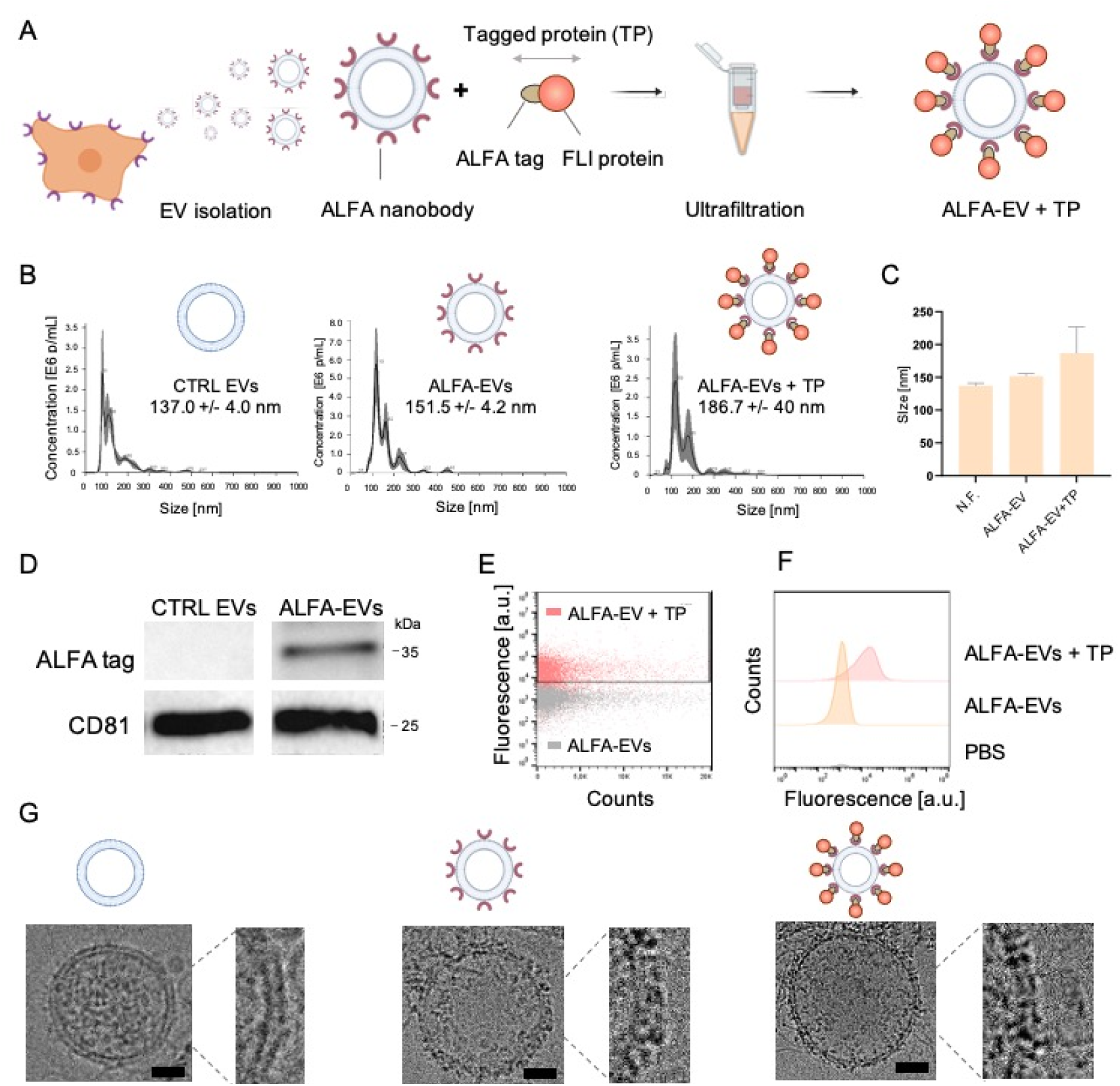
Characterization of ALFA nanobody EVs. **(A)** EVs were isolated from parental ALFA nanobody-displaying cells, incubated with ALFA-tagged proteins, and purified by ultrafiltration creating functionalized ALFA-EVs. **(B)** Nanoparticle tracking analysis (NTA) of the formulations revealed increasing size with binding of tagged proteins on EV surface. **(C)** Quantification of data in B. **(D)** Western blot analysis of protein expression in isolated EVs showed exclusive presence of the nanobodies on ALFA-EVs. **(E)** Nano flow cytometry dot plots of EVs with clicked dTomatoTP proteins (ALFA-EV + TP) and controls without fluorescent proteins (ALFA-EV). **(F)** Corresponding histograms to E. **(G)** Cryo-electron microscopy images of non-functionalized EVs (left), ALFA-EVs (middle) and ALFA-EVs with clicked mNeonTP proteins (right). The insets show a detailed view of the membrane bound proteins. Scale bar is 30 nm.

NTA showed that ALFA-EVs and functionalized ALFA-EVs were both slightly larger than control EVs, and functionalized ALFA-EVs were larger than non-functionalized ALFA-EVs (187±40, 151±4, and 137±4 nm, respectively; **Fig. 2B, C)**. Cryo-EM analysis of EVs revealed a similar unilamellar structure with a pronounced lipid bilayer in all EV formulations (control, ALFA-EVs and functionalized ALFA-EVs) (**Fig. 2G, Fig. S6**), and ALFA-EVs revealed compact hypointense spots on the surface of the EVs, suggesting the ALFA nanobodies were present at high density (**Fig. 2G insets**).

Finally, Flow Induced Dispersion Analysis (FIDA) showed that ALFA-EVs bind strongly and specifically to ALFA-tagged proteins under physiological flow conditions (**Fig. 3A**). This multimodal analysis concluded that ALFA-EVs maintain the properties of intact EVs and that ALFA-tagged proteins specifically bind to ALFA-EVs. Finally, we evaluated the long-term stability of functionalized ALFA-EVs. Nanoflow cytometry revealed that the ALFA nanobody is stable on the surface of EVs over a period of 5 months (**Fig. S8**). Although the ALFA-tagged proteins dissociate over this period, ALFA-EVs can easily be re-functionalized even after 5 months (**Fig. S8**).

**Fig. 3.**
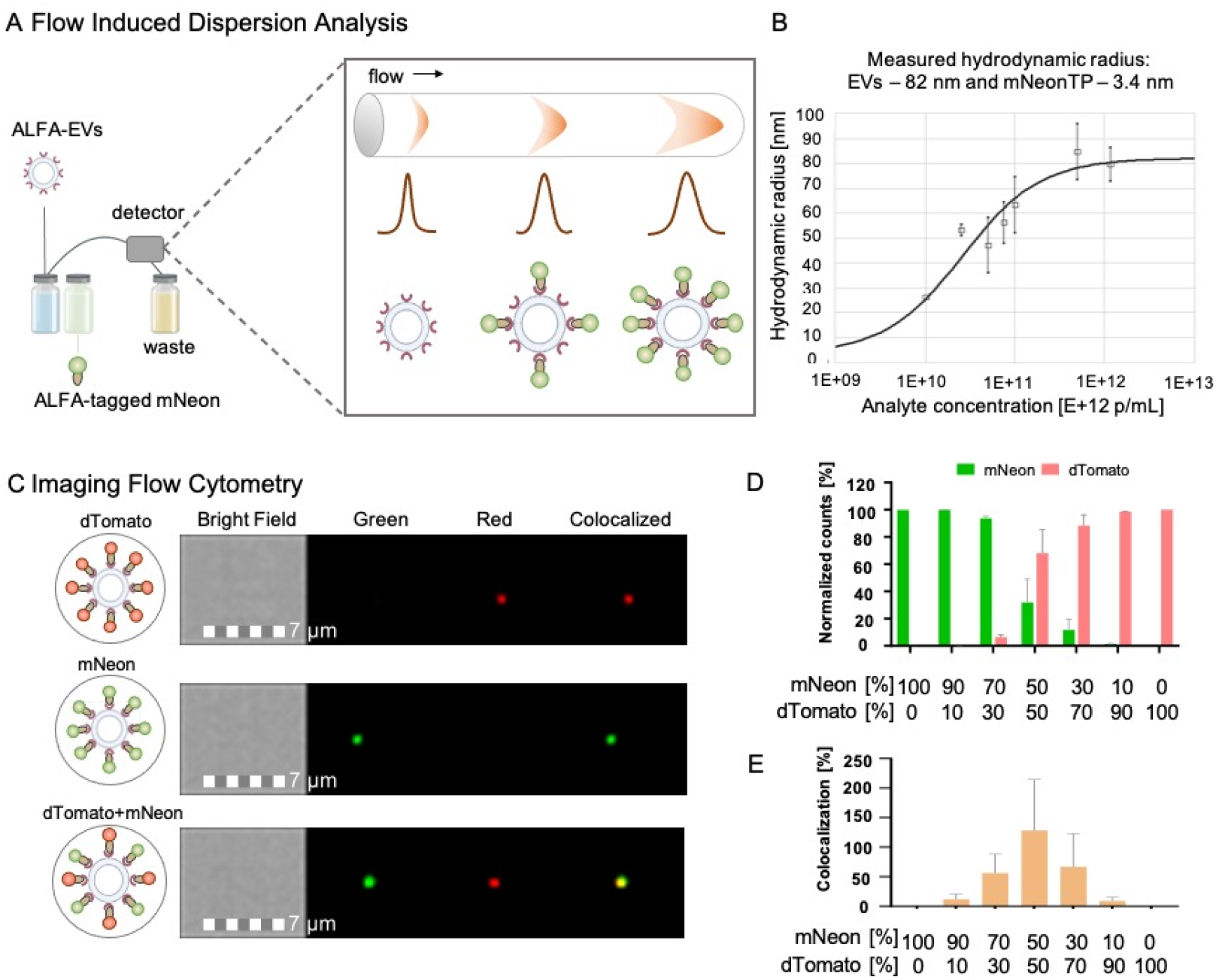
Functionalization of ALFA-EVs. **(A)** An experimental scheme of the Flow Induced Dispersion Analysis (FIDA). Briefly, the ALFA-tagged protein is mixed with EVs in a capillary under continuous flow, and the changes in fluorescent signal intensities reflecting the diameter of the particles are being recorded. **(B)** The binding of ALFA-tagged mNeon to ALFA-EVs at various concentration of EVs measured with FIDA. FIDA revealed the hydrodynamic radius of the vesicles to be 82 nm and the bound proteins 3.4 nm **(C)** Imaging Flow Cytometry images of single ALFA-EVs with clicked dTomatoTP and mNeonTP. **(D)** Quantification of fluorescent signals from EVs functionalized at different ratios of dTomatoTP:mNeonTP proteins. **(E)** Quantification of co-localization of dTomatoTP and mNeonTP signals.

### 2.3 Simultaneous functionalization of EVs with multiple proteins

We next examined whether NaTaLi can functionalize EVs with multiple proteins fused to the ALFA tag at the same time. ALFA-EVs were functionalized with mNeonTP and dTomatoTP and multispectral imaging flow cytometry (ImageStream) confirmed distinct fluorescent signals corresponding to each protein. Upon simultaneous incubation with both proteins, co-localization of the two fluorescent signals was observed, indicating multiple display on individual EVs (**Fig. 3C, Fig. S9**). Moreover, ALFA-EVs were incubated with mNeonTP and dTomatoTP proteins at different concentration ratios and higher mNeon/dTomato concentration ratios led to higher mNeon/dTomato signal on the EV surface (**Fig. 3D, E**). We thus concluded that NaTaLi enables ratiometric and “one-pot” functionalization of EVs with multiple proteins.

### 2.4 Targeting of EVs to cancer cells

Having established that ALFA-EVs can robustly be functionalized with ALFA-tagged proteins, we next evaluated their potential for *in vitro* and *in vivo* cancer cell targeting. ALFA-EVs were functionalized with the tumor-targeting peptides RGD and LinTT1 (**Fig. 4A**). RGD peptide binds to integrins overexpressed on cancer cells^23^, whereas LinTT1 peptide targets the p32 protein overexpressed by breast cancer and cancer-associated cells^24^. Functionalization with either peptide led to significantly increased uptake of functionalized EVs in cancer 4T1 cells compared to control EVs, without a significant increase in uptake in non-cancerous HEK293 cells (**Fig. 4C, D**).

**Fig. 4.**
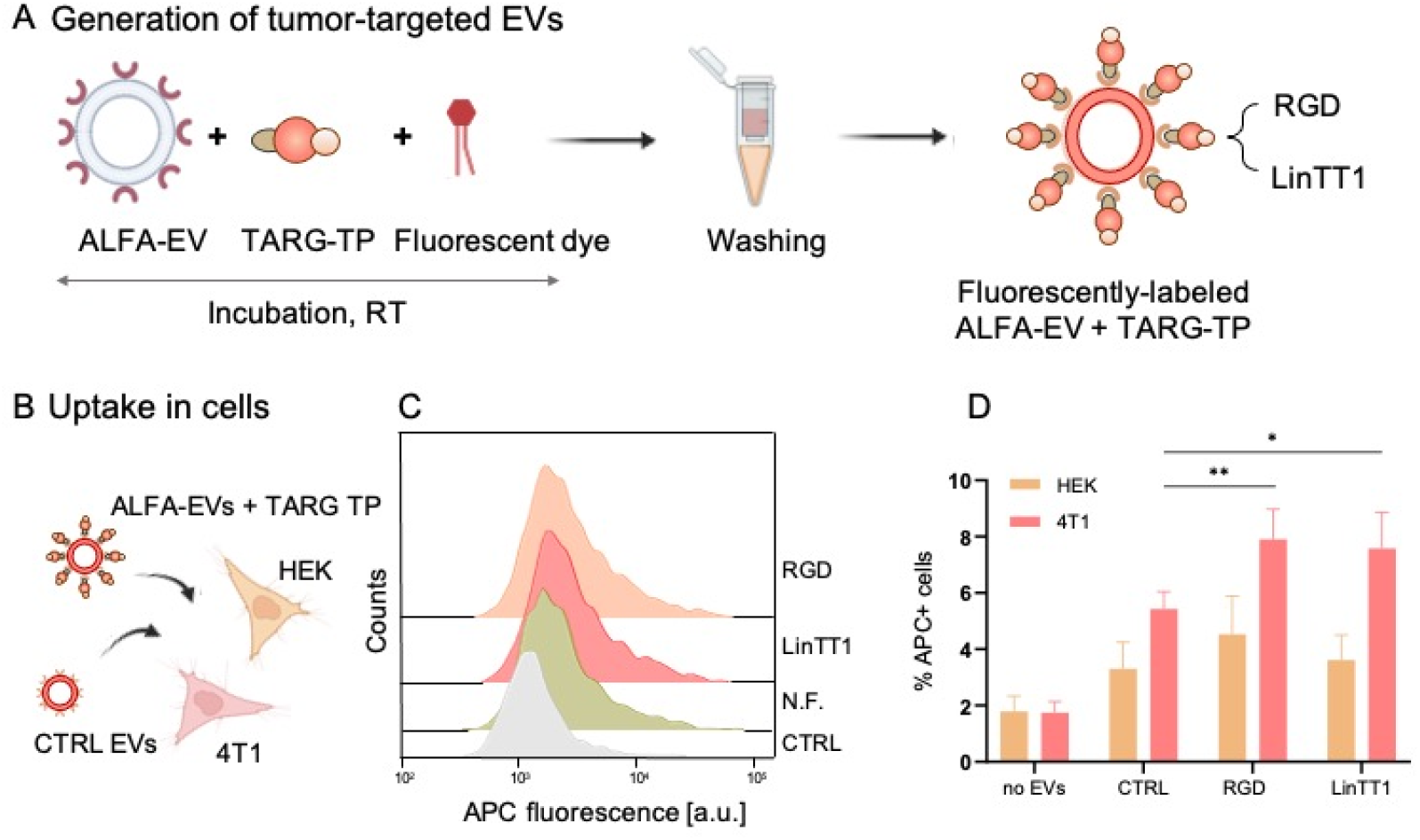
Tumor targeting by ALFA-EVs. **(A)** EVs were functionalized with dTomato-fused targeting peptides RGD or LinTT1 and a fluorescent. **(B)** A scheme of the uptake experiment in cancer (4T1) and benign (HEK293) cells. **(C)** Flow cytometry histograms of 4T1 breast cancer cells with accumulated tumor-targeted and control EVs. **(D)** Quantification of the uptake of EVs in HEK293 and cancer 4T1 cells.

Next, the biodistribution and targeting of ALFA-EVs functionalized with RGD or LinTT1 peptides were tested in a mouse model of breast cancer *in vivo*. Because dTomato fluorescence overlaps with the autofluorescence of tissues (**Fig. S10G**), EVs were incubated with the near-infrared fluorescent dye DiR before injection, allowing a distinct fluorescent signal outside of the range of tissue autofluorescence. Tumor-targeted functionalized EVs showed both dTomato and DiR fluorescence signal, while the non-functionalized EVs lacked the dTomato signal (**Fig. S10B, C**). After intravenous administration, EVs were detected in the subcutaneous tumors in mice, with RGD- and LinTT1-functionalized EVs leading to significantly higher accumulation in tumors compared to control ALFA-EVs (**Fig. 5B**) despite similar signal in the liver (**Fig. 4A**).

**Fig. 5.**
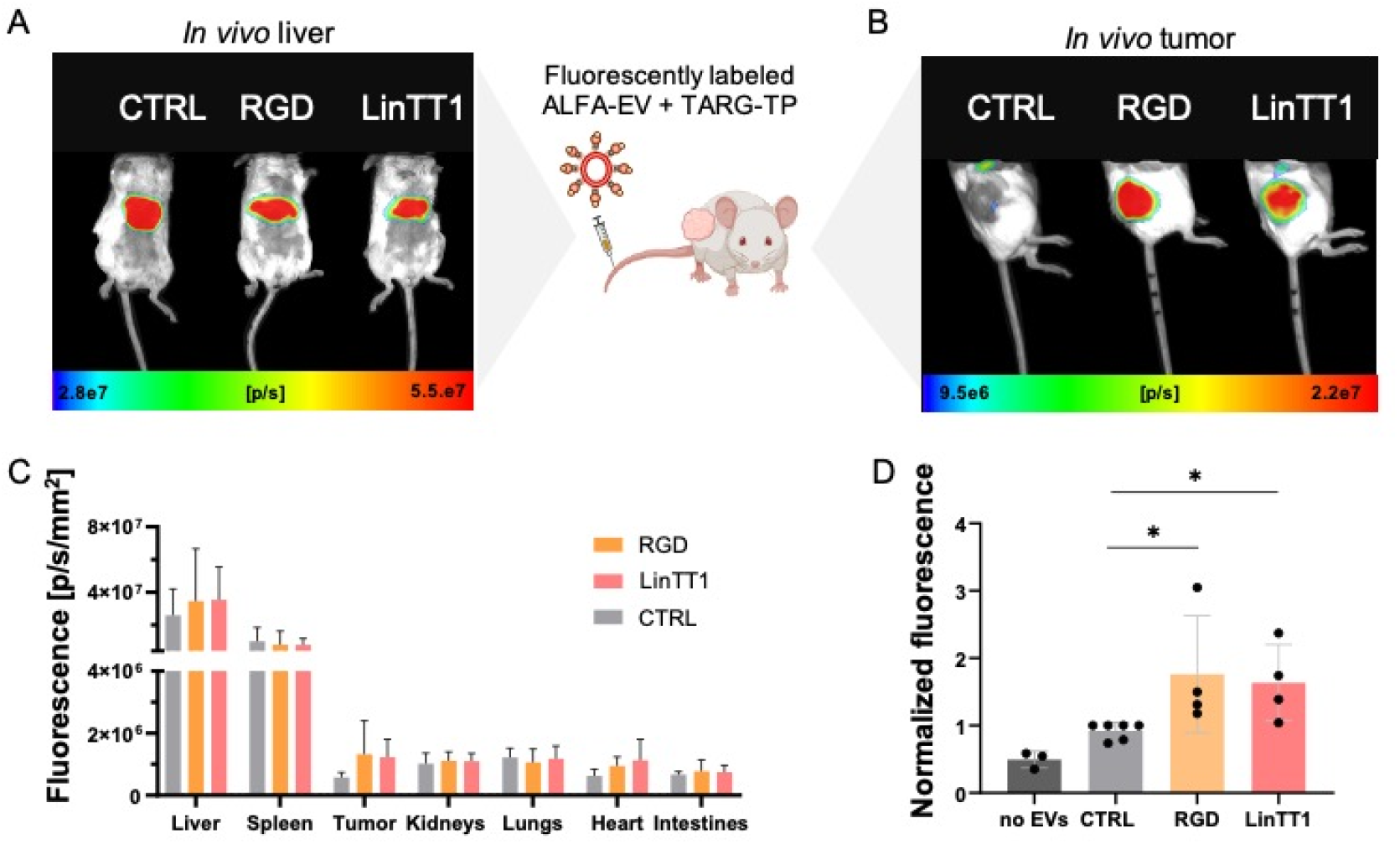
*In vivo* targeting and biodistribution of tumor-targeted NaTaLi EVs. **(A)** *In vivo* fluorescence imaging shows similar accumulation of functionalized and non-functionalized EVs in mouse livers at the time point 24 h after intravenous administration. **(B)** Significantly higher uptake or tumor-targeted RGD- and LinTT1-functionalized EVs in tumors at the time point 24 h after i.v. injection. **(C)** Biodistribution of the functionalized and control EVs in various organs at 24 hours after EV administration. **(D)** Quantification of the *in vivo* fluorescence signals emitted from the tumors showed significant enhancement of accumulation by functionalized EVs. Fluorescence signals of RGD-and LinTT1-EVs in tumors are normalized to fluorescence of control EVs.

This result was supported by *ex vivo* analysis of harvested organs (**Fig. 5E, F**), which revealed enhanced tumor accumulation of functionalized EVs and similar accumulation of functionalized and non-functionalized EVs in the liver, spleen, kidneys, intestines, lungs and hearts (**Fig. 5E**). These results suggest that functionalized ALFA-EVs exhibit tumor targeting without substantial other changes in biodistribution, confirming the NaTaLi approach can robustly functionalize EVs for *in vivo* targeting.

## 3. Discussion

EVs can be functionalized through a wide range of strategies that improve their targeting, stability, and therapeutic performance^25–27^. Current approaches for chemically functionalizing EVs or inserting proteins into the membrane of EVs can lead to degradation of EV integrity. In contrast, genetically engineering parental cells to express membrane-anchored fusion proteins allows for robust display of versatile proteins without membrane disruption. Conventional approaches for EV genetic engineering rely on generating a separate stable cell line for each individual surface protein and are therefore time and labor-intensive^7,10,27,28^. Moreover, proteins can express and be displayed at very different levels, leading to substantial variability. These issues significantly slow the development of EV-based drug delivery systems and limit their broader application in translational medicine.

To address these challenges, we developed the NaTaLi system, in which protein-tag-binding nanobodies are displayed on the surface of EVs. Once a single stable cell line is established to create nanobody-displaying EVs, any protein or peptide fused to the protein tag can be easily displayed on the surface of EVs. Moreover, functionalization of EVs using NaTaLi relies only on the expression and display of the protein-tag-binding ALFA nanobody and the affinity of the nanobody to the protein ALFA tag, which are both constant. Therefore, in addition to reducing the time and effort needed to functionalize EVs, NaTaLi likely results in substantially lower variability in display levels between different proteins than other approaches. Another important aspect of the NaTaLi system is that the targeting peptides are produced recombinantly in bacteria, dramatically reducing the amount of work with mammalian cell cultures and significantly accelerating development cycles.

The ALFA nanobody represents a universal docking interface and an excellent targeting system for EVs due to its high binding affinity to the ALFA tag. We proved that the ALFA nanobody-ALFA tag affinity is in picomolar range with very low dissociation effect in our NaTaLi system. Strong binding of ALFA-EVs to ALFA-tagged proteins was shown also under the *in vivo* mimicking conditions which confirms an excellent stability and versatility of the NaTaLi system for *in vivo* applications. ALFA nanobody can be also readily fused with to virtually any protein^19^. We showcased functionalization of ALFA-EVs with various fluorescent proteins and tumor-targeting peptides.

Crucially, NaTaLi enabled us to test several different peptides for cancer-cell targeting through producing only one stable cell line. Thus, NaTaLi enables much higher-throughput testing of EV functionalization than other approaches. Moreover, we demonstrated that NaTaLi allows ratiometric functionalization of EVs with multiple different proteins at the same time in “one pot”. Recent work has shown that display of multiple proteins on the surface of EVs can lead to enhanced cellular uptake of EVs *in vitro*^25,29,30^ and NaTaLi provides a streamlined and robust approach to multiplex display.

Finally, we demonstrated that NaTaLi-functionalized EVs carrying tumor-targeting peptides efficiently accumulated in tumors *in vivo* while maintaining biodistribution profiles similar to control EVs. Their ability to robustly distinguish diseased tissue illustrates the potential of the NaTaLi platform for targeted delivery of therapeutic cargos. Achieving selective tumor accumulation is a major goal in the development of next-generation delivery systems^31^. Although liposomes^32^, synthetic nanoparticles^33^, polymers^34^, polymerosomes^35^ and other materials have been explored, EVs offer distinct advantages including superior biocompatibility, low immunogenicity, and robust functionalization capacity. The ease of use and modularity of NaTaLi positions it as a promising system for the future development of precision EV-based therapies.

## 4. Conclusion

We demonstrate the development of the Nanobody-Targeting-Ligand (NaTaLi) system, a highly versatile platform for multiplexed functionalization and precise targeting of extracellular vesicles. Using advanced analytical technologies, we confirmed strong and highly specific interactions between displayed nanobodies and their complementary targeting peptides on both engineered cells and EVs. *In vivo* studies in tumor-bearing mice further validated the system, revealing clear accumulation of targeted vesicles within tumor tissue. Together, these findings establish NaTaLi as a robust and adaptable strategy that opens new avenues for multiplexed cancer biomarker detection and the targeted delivery of therapeutics by extracellular vesicles.

## 5. Material and methods

### Cloning and DNA manipulation

The sequence for the designed ALFA nanobody (DnbALFA^18^) (**Fig. S1**) was cloned into the pAGD plasmid^28^ using the Restriction-free cloning (RFC)^36^. The final pAGD_ALFA plasmid contains CMV promoter, the resistance to puromycin that is fused with eUnaG2 fluorescent protein at the C-terminus (expression in cytoplasm) and ALFA nanobody (extracellular expression) fused with the platelet-derived growth factor receptors (PDGFRβ) transmembrane domain.

ALFA tag sequence (SRLEEELRRRLTE) was designed as fusion with a protein of choice and cloned into pET28-bdSUMO expression plasmid^37^ by RFC. The ALFA tag and Strep tag (WSHPQFEK) fused with bdSUMO sequence with His tag to allow specific purification by StrepTactin and Ni-NTA Sepharose and bdSUMO protease (**Fig. S1**). A sequence for a bright monomeric green fluorescent protein mNeon^38^ and red fluorescent protein dTomato^39^ were fused between the His tag and ALDA tag. A sequence of a targeting peptide RGD (CRGDKGPDC^23^) and LinTT1 (AKRGARSTA^24^) was cloned into the pET28-bdSUMO plasmid between the ALFA tag and the Strep tag sequence. An NGL linker (SGGGGSGGGGNGSNGSGGSS) was cloned between the ALFA tag and the targeting peptide to provide a distance between the peptide and the phospholipid membrane. SPY tag (AHIVMVDAYKPTK) was used as a control tag for the tagged proteins (ctrlTP).

### Preparation of ALFA-tagged proteins

ALFA-tagged proteins (TP) were purified from *E*.*Coli* BL21 (DE3) as already described^28^. Briefly, *E*.*Coli* were transformed with a pET_bDSUMO plasmid described above (10 ng of plasmid). Then, 200 mL of 2YT medium (16 g of tryptone, 10 g of yeast extract, and 5 g of NaCl, pH 7) was inoculated (1%), grown to the OD600 = 0.6 at 37 °C. The expression was initiated by the addition of IPTG to a final concentration of 0.5 mM. The expression continued for the next 16 h at 30 °C. Expressed cells were washed (50 mM Tris-HCl, 200 mM NaCl buffer, pH 8) and disintegrated by sonication (40 W, 20s pulse length, 40s pause). Subsequently, the cell debris was pelleted by 6000g, 10 mins and the supernatant was purified using Ni-NTA agarose (PureCube Ni-NTA agarose, Cube Biotech, Germany) followed by StrepTactin Sepharose (Cube Biotech, Germany). The eluted fraction was further purified by size exclusion chromatography on HiLoad 26/600 Superdex 75 using NGS 10 size-exclusion system (Biorad, USA). Four different proteins were isolated nad purified: ALFA-tagged mNeon (mNeonTP), ALFA-tagged dTomato (dTomatoTP), ALFA-tagged dTomato-RGD (dTomatoTP+RGD) or ALFA-tagged dTomato-LinTT1 (dTomatoTP+LinTT1).

### Cell culture

All cell lines were cultured at 37 °C in a 5% CO_2_ atmosphere and regularly screened for mycoplasma contamination. The following cell lines were used: Human embryotic kidney cells (HEK293; ATCC No.), mouse breast cancer cells (4T1; ATCC No. CRL-2539), bone metastasis adenocarcinoma cells (PC3; ATCC No. 1435) and Chinese hamster ovarian cancer cells (CHO; ATCC No. CCL-661).

### Preparation of ALFA nanobody expressing cells

Human embryotic kidney cells (HEK293) were cultured in DMEM high glucose (4.5 g/L) medium supplemented with 10% fetal bovine serum (FBS) and 1% penicillin/streptomycin antibiotics. Cells were passaged 2–3 times per week using trypsin EDTA solution A (Sigma Aldrich, USA) for cell detachment. For transfection, the cells were seeded a day before transfection at the confluency of 30%. A mixture of a transfection reagent (jetPEI, Polyplus, France) and a plasmid was added to the cells according to the manufacturer’s recommendations and incubated for 24 or 48 h. The stable cell line was established after fluorescent activated cell sorting (FACS) and long-term exposure of cells to selection antibiotics (1 µg/mL puromycin) in 4–6 weeks.

### Flow cytometry of ALFA nanobody-displaying cells

The cells were cultured until full confluence, then they were detached from the plates by PBS and added to the 1.5 mL Eppendorf tubes. After spinning down (5 min at 500 g), the cell pellet was resuspended in a labeling solution containing fluorescently labeled, purified tagged protein in PBS supplemented with 2% FBS. The cells were labeled for 1 h on ice and then washed two times with 1 mL of PBS (spun down each time for 5 min at 500 g). After the last wash, the cells were resuspended in PBS with 2% FBS solution and added to the FACS tubes. The fluorescence was measured by the Navios Galios FACS machine (Beckman Coulter, USA) and analyzed by the FlowJo software.

### Ligand Tracer

ALFA nanobody-displaying and control (non-functionalized) HEK293 cells were seeded in a MultiDish 2×2 according to the manufacturer’s instructions. Briefly, the plate was coated by incubation for 45 min with fibronectin (0.01 mg/mL of PBS) (Sigma Aldrich, USA) to enhance the cell attachment. Then, cell suspension (1 mil) was pipetted in a 1 cm drop and kept incubating for 45 min. Unattached cells were removed and the plate was filled with DMEM culture medium with 10% FBS. For measurement, medium was supplemented with various concentration of mNeonTP (10, 50 and 150 nM) and the binding was monitored in real time by the Ligand Tracer Green 2nd gen (Ridgeview Instruments, Sweden). The instrument was placed on a bench with temperature regulated to 25 °C. MultiDish 2×2 with cells was placed in the instrument with immobilized positive and negative cells. One cycle was set for 18 seconds detection per spot, 3 s delay, 15 s signal uptake. The rotating platform periodically circulated with pure DMEM medium for baseline establishment. Once the baseline was established, the 10 nM mNeonTP protein was introduced to the medium. The signal was recorded until equilibrium state was achieved with defined concentration. After equilibrium state was achieved, 50 nM concentration was introduced to the system, etc for 150 nM. After reaching equilibrium state for 150 nM protein, the measurement was stopped, medium was replaced with fresh DMEM medium, and the dissociation was measured for 3 hours to receive 10 % decrease in signal for accurate dissociation constant characterization. Recorded binding curves were evaluated with TraceDrawer software (Ridgeview Instruments, Sweden) following single cycle kinetics model.

### Binding of ALFA-tagged proteins to ALFA-nanobody cells

ALFA nanobody-expressing HEK293 cells were cultured under regular conditions in full DMEM medium (10% FBS) and 1% penicillin/streptomycin antibiotics) supplemented with 1 μg/mL puromycin, the control non-functionalized cells were cultured in the same medium without puromycin. The ALFA-expressing and control HEK293 cells were seeded in a 6-well plate. Next day, mNeonTP or dTomatoTP proteins were added to the cells at concentration of 100 nM and incubated for 1h at 37 °C. The cells were then washed twice with PBS and fresh phenol-free DMEM medium (Sigma Aldrich, USA) was added to the cells. Cells were examined under a fluorescence microscope APX100-HCU (Evident Scientific, Japan) using specific filters for mNeon (exc/em = 470/508 nm) and dTomato (exc/em = 545/580 nm).

### Binding of cancer-targeting ALFA-tagged proteins to cells

Cancer cells (CHO, PC3, 4T1) were cultured at 37 °C in a 5% CO_2_ atmosphere in DMEM (PC3 cells) or RPMI (4T1 and CHO cells) medium supplemented with 10% fetal bovine serum (FBS) and 1% penicillin/streptomycin antibiotics. For the experiment, the cells were detached with PBS, spin down at 200 g for 5 min and places into 1.5 mL tubes. Isolated and purified ALFA-tagged and cancer targeted proteins (dTomatoTP+LinTT1, dTomatoTP+RGD) of concentration 200 nM were incubated with cancer cells (500k cells/tube) for 1h at RT and regularly vortexed. Then the cells were washed two times with PBS (spin 200 g, 5 min) and resuspended in the FACS buffer (PBS with 2% FBS). Cells were examined by the Navios Galios FACS machine (Beckman Coulter, USA) and analyzed by the FlowJo software. The normalized fluorescence was expressed as % of the fluorescence signal of the non-labeled cells.

### Western blot

For the Western blot analysis, EVs or parental cells were lysed by RIPA buffer (1× for cells, 1:1 in PBS for EVs) supplemented with a proteinase inhibitor cocktail. Proteins were separated on ExpressPlus PAGE Ready gels 4–20% (A2S, M42015) and transferred on the cellulose membrane (300 mA, 90 min) using a BioRad blotting device (BioRad, USA). After overnight blocking in 5% milk in TBST, specific antibodies were applied to the membranes for 1 h to detect markers: CD81 (B–II, Santa Cruz, USA); c-myc (9E10, Santa Cruz, USA) and beta-actin (Sigma Aldrich USA) (1:1000). HRP-conjugated antimouse secondary HRP goat antimouse IgG antibody (No. 4053, Biolegend, USA) (1:5000 in TBST) was applied for 1 h before imaging using enhanced chemiluminescence substrate EZ-ECL Kit (ThermoFisher, USA).

### Isolation of ALFA nanobody-expressing EVs

EVs were isolated from ALFA nanobody-expressing HEK293 or non-functionalized control HEK293 cells using differential ultracentrifugation methods combined with ultrafiltration (100 kDa)^28^. Briefly, cells were cultured in DMEM supplemented with 10% exosome-free serum (Gibco, USA) and 1% penicillin/streptomycin antibiotic solution for 48 – 72 hours. Then, the culture medium from the incubating cells was collected and centrifuged at 400 g for 10 min, 2000 g for 10 min and 10,000 g for 30 min at 4 °C for the elimination of cells, cell debris, microvesicles and apoptotic bodies. The cleared medium was then concentrated using 100 kDa JumboSep filters (Pall Corporation, USA) by centrifugation at 3000 g for 30 min). Finally, the EVs were pelleted at 100,000 g using the Beckman Sw40 rotor (Beckman Coulter, USA) at 4 °C for 2 hours. Then the pellet was resuspended in PBS, the tubes were filled with PBS and the final EV pellet was spun down at 100,000 g for 2 hours. The pellet was then resuspended in 0.2 μm-filtered PBS and stored at −80 °C until further use.

### Nanoparticle tracking analysis

The size and concentration of isolated EVs was measured by nanoparticle tracking system (NanoSight NS300, Malvern Instruments, UK) using with a 405 nm laser by acquiring three 1 min videos at the camera level 16. Threshold 5 was used for the analysis of all samples.

### Cryogenic Electron Microscopy (Cryo-EM)

4 µl of EV sample was applied to freshly plasma-cleaned TEM grids (Quantifoil, Cu, 300mesh, R2/1) and vitrified into liquid ethane using ThermoScientific Vitrobot Mark IV (5°C, 100% rel. humidity, 30 s waiting time, 6 s blotting time). The grids were subsequently mounted into the Autogrid catridges and loaded to Glacios 2 (ThermoScientific) transmission electron microscope for imaging. The microscope was operated at 200kV. The exosome cryo-TEM micrographs were collected on Falcon4i direct electron detection camera at the 11,000x and 190,000x nominal magnification with the underfocus in the range 2–4 µm and the overall dose of ∼20 e/Å2.

### Preparation of functionalized EVs

Isolated EVs were resuspended with 100 nM of ALFA-tagged proteins (mNeonTP, dTomatoTP, dTomatoTP + RGD, dTomatoTP + LinTT1) and incubated for 1 hour at room temperature while shaking. Then the unbound proteins were washed by ultrafiltration using Amicon filters (100 kDa, 0.5 mL) by centrifugation at 14,000g for 5 mins (three times) and 1000g for 2 mins (reverse position). Final EV solution was resuspended in 0.2 µm filtered PBS.

### Nano flow cytometry

EVs tagged with fluorescent proteins were measured with a nano flow cytometer Explorer 3 (VdioBiotech, China). EVs were diluted to concentration of 1×10^9^ per mL in PBS. EVs were measured with 488 nm excitation and 525 emission filter. PBS used for dilution of the samples was used as a control. Fluorescently-functionalized EVs were measured immediately after tagging and 5 months after tagging with mNeonTP and re-tagged after 5 months storage in the freezer at −20 °C.

### Flow induced Dispersion Analysis

Binding of tagged proteins to ALFA nanobodies on EVs was characterized by Flow Induced Dispersion Analysis (FIDA Biosystems Aps, DK)^40^. For this, ALFA nanobody EVs were freshly isolated from ALFA nanobody-expressing cells. EVs of different concentration (ranging from 1×10^8^ to 2×10^12^) were supplemented with 1% pluronic acid F127 (Sigma Aldrich, USA) to prevent sticking to the capillary. EVs were placed into the experimental vials with inserts. Another vial was filled with 80 nM mNeonTP and PBS as a control. Fluorescence was monitored over time. FIDA method applied for this measurement is as follows: 1M NaOH rinsing 45s under 3500 mBar, MiliQ washing 60 s under 3500 mBar, analyte injection 30 s under 3500 mBar, indicator injection 10 s under 50 mBar, diffusivity acquisition 1800 s under 50 mBar. The analyte for this measurement was either buffer or corresponding concentration of EVs. The indicator for this measurement was 80 nM mNeonTP protein in PBS. Capillary mixing was used for this measurement. As is indicator injected to preequilibrated capillary filled with analyte it immediately starts binding to the analyte (or does not bind in case of buffer). The second approach is used for the indicator screening to receive information about the size of the indicator and also about the viscosity of the system. Once the buffer is replaced with corresponding EVs concentration the binding is observed accordingly. Binding curve was evaluated with FIDA Software V3.0 Beta (FIDA Biosystems Aps, Denmark).

### Imaging flow cytometry

ALFA nanobody EVs were functionalized with mNeonTP and dTomatoTP at concentration of 80 nM. Double functionalized EVs were incubated with proteins at different ratio of mNeonTP:dTomatoTP, 100 nM:0 nM; 70 nM:30 nM; 50 nM:50 nM; 30 mM:70 nM; 0 nM:100 nM. Unlabeled EVs and PBS with 80 nM of the proteins were used as controls. EVs were measured using a multispectral ImageStreamX Mark II imaging flow cytometer (Amnis Corp., part of Cytek, CA) using a 60× lens (NA = 0.9). The lasers used were 488 nm (400 mW) and 561 nm (200 mW), and the channels collected were bright field (Ch01), green (Ch02) and red (Ch04). At least 20□000 events were recorded from each sample. Data was analyzed using the manufacturer’s image analysis software (IDEAS 6.3; Amnis Corp.). Objects were gated for low brightfield area and low side scatter values, to exclude debris and the speedbeads that run with sample as part of instrument operation. The fluorescent signal was quantified using the Max pixel values of channel 2 (for mNeon) and channel 4 (for dTomato). Gating was set according to the unstained controls.

### *In vitro* binging of targeted EVs to cells

EVs with clicked ALFA-tagged cancer targeted proteins (RGD and LinTT1) were labeled by ExoBrite dye according to the manufacturer’s instructions (incubation 1h at RT). Then the unbound dye was removed by ultrafiltration using Amicon filters (100 kDa, 0.5 mL) by centrifugation at 14,000 g for 5 mins (three times) and 1000g for 2 mins (reverse position). EVs were then incubated with cancer cells (4T1, PC3, CHO) for 2 hours at 37 °C (1×10^9^ EVs/mL). After two washes, the cells were detached with trypsin and examined by the Navios Galios FACS machine (Beckman Coulter, USA). The FACS data were analyzed by the FlowJo software.

### Animal model

Balb/c mice (Velaz, Germany) were used in the experiments. All animal studies were approved in accordance with the European Communities Council Directive (2010/63/EU) and the national laws (ethical approval No. 50/2022). All animals were kept in a daily controlled room at the Institute for Clinical and Experimental Medicine animal facility with a surrounding relative humidity level of 50 ± 10% and a temperature of 22 ± 1 °C, with a 12/12 cycle of dark and light phases. Subcutaneous xenograft tumors were induced by injection of 4T1 mouse breast cancer cells under the skin above the mouse flank. Mice were kept under isoflurane anesthesia during the whole procedure. Each mouse received 3 mil of cells 100 μL of PBS. The tumors were allowed to grow for 2–3 weeks until they reached a sufficient size (diameter around 0.5 – 1 cm).

### *In vivo* fluorescence imaging

For *in vivo* experiments, cancer targeted ALFA-tagged EVs were labeled with DiR fluorescent dye (ThermoFischer Scientific, D12731) at a concentration of 5 μM by incubation for 1 h at room temperature. EVs were then washed three times with the 100 kDa Amicon centrifugation filters (5 min at 14,000g, and 2 min at 1000g in a reverse position). Then, mice with subcutaneous tumors were injected with 100 μL solution of 1×10^11^ tumor-targeted ALFA-tagged EVs retroorbitally. Two types of tumor-targeted EVs (LinTT1 and RGD) were administrated. Control ALFA-EVs (CTRL) were prepared in the same way but without incubation with the cancer-targeted proteins. Control animals (CTRL) received injection of 100 μL of PBS. Mice were measured with an optical imager Ami HT (Spectral Instruments, Bruker, Germany) using 5 s exposure time, 50 % power and excitation and emission filters for DiR (750/790 nm). After the last time point (24 hours), the mice were sacrificed and the individual organs (liver, spleen, kidneys, heart, lungs, intestine) were harvested and fixed with 4% formaldehyde solution in PBS. All organs were visualized using the optical imager while keeping the same parameters as for the *in vivo* measurements.

### Statistical analysis

All numerical data are presented as mean ± standard deviation (S.D.). Statistical analysis was performed using GraphPad Prism 8.0 software (GraphPad Software Inc., USA). Comparison of two groups was analyzed by a two-tailed Student’s t test. A p-value of 0.05 and below was considered significant: *p-value <0.05, **p-value <0.01, ***p-value <0.001, and ****p-value <0.0001.

## Acknowledgement

This work was supported by the Czech Science Foundation (GACR) - Grant No. 22-13334I and the Ministry of Health CR-DRO (Institute for Clinical and Experimental Medicine IKEM, IN00023001).

We also acknowledge the CIISB, Instruct-CZ Centre, supported by MEYS CR (LM2023042) for acquisition of the cryo-EM images and the Light Microscopy Core Facility, IMG ASCR, Prague, Czech Republic, supported by MEYS - LM2023050, MEYS - CZ.02.1.01/0.0/0.0/18_046/0016045 and MEYS - CZ.02.01.01/00/23_015/0008205, for their support with the ImageStream imaging presented herein.

## Author contributions

**Andrea Galisova:** Conceptualization, performing biological assays and in vivo experiments, data analysis, supervision, writing and editing. **Jiri Zahradnik:** Conceptualization, protein design. **Ester Merunkova:** performing biological assays. **Dominik Havlicek:** performing in vivo experiments and data analysis. **Josef Uskoba:** performing FIDA and Ligand Tracer experiments and data analysis. **Ziv Porat:** analysis of ImageStream data. **Daniel Jirak:** supervision, writing and editing.

## References

1. Kalluri R, McAndrews KM. The role of extracellular vesicles in cancer. Cell. 2023;186(8):1610–1626. doi:10.1016/j.cell.2023.03.010

2. Couch Y, Buzàs EI, Di Vizio D, et al. A brief history of nearly EV-erything - The rise and rise of extracellular vesicles. J Extracell vesicles. 2021;10(14):e12144. doi:10.1002/jev2.12144

3. Teng F, Fussenegger M. Shedding Light on Extracellular Vesicle Biogenesis and Bioengineering. Adv Sci (Weinheim, Baden-Wurttemberg, Ger. 2020;8(1):2003505. doi:10.1002/advs.202003505

4. Herrmann I, Wood M, Fuhrmann G. Extracellular vesicles as a next-generation drug delivery platform. Nat Nanotechnol. 2021;16:748–759.

5. Wiklander OPB, Nordin JZ, O’Loughlin A, et al. Extracellular vesicle in vivo biodistribution is determined by cell source, route of administration and targeting. J Extracell vesicles. 2015;4:26316. doi:10.3402/jev.v4.26316

6. O’Brien K, Breyne K, Ughetto S, Laurent LC, Breakefield XO. RNA delivery by extracellular vesicles in mammalian cells and its applications. Nat Rev Mol Cell Biol. 2020;21(10):585–606. doi:10.1038/s41580-020-0251-y

7. Ohno S, Takanashi M, Sudo K, et al. Systemically injected exosomes targeted to EGFR deliver antitumor microRNA to breast cancer cells. Mol Ther. 2013;21(1):185–191.

8. Alvarez-Erviti L, Seow Y, Yin H, Betts C, Lakhal S, Wood MJA. Delivery of siRNA to the mouse brain by systemic injection of targeted exosomes. Nat Biotechnol. 2011;29(4):341–345. doi:10.1038/nbt.1807

9. Wiklander OPB, Mamand DR, Mohammad DK, et al. Antibody-displaying extracellular vesicles for targeted cancer therapy. Nat Biomed Eng. Published online 2024. doi:10.1038/s41551-024-01214-6

10. El-Andaloussi S, Lee Y, Lakhal-Littleton S, et al. Exosome-mediated delivery of siRNA in vitro and in vivo. Nat Protoc. 2012;7(12):2112–2126.

11. Usman WM, Pham TC, Kwok YY, et al. Efficient RNA drug delivery using red blood cell extracellular vesicles. Nat Commun. 2018;9(1):2359. doi:10.1038/s41467-018-04791-8

12. Berggreen AH, Petersen JL, Lin L, Benabdellah K, Luo Y. CRISPR delivery with extracellular vesicles: Promises and challenges. J Extracell Biol. 2023;2(9):e111. doi:10.1002/jex2.111

13. Smyth T, Petrova K, Payton N, et al. Surface Functionalization of Exosomes Using Click Chemistry. Bioconjug Chem. 2014;25(10):1777–1784.

14. Gao X, Ran N, Dong X, et al. Anchor peptide captures, targets, and loads exosomes of diverse origins for diagnostics and therapy. Sci Transl Med. 2018;10(444):eaat0195. doi:10.1126/scitranslmed.aat0195

15. Haraszti RA, Miller R, Stoppato M, et al. Exosomes Produced from 3D Cultures of MSCs by Tangential Flow Filtration Show Higher Yield and Improved Activity. Mol Ther. 2018;26(12):2838–2847. doi:10.1016/j.ymthe.2018.09.015

16. Ivanova A, Badertscher L, O’Driscoll G, et al. Creating Designer Engineered Extracellular Vesicles for Diverse Ligand Display, Target Recognition, and Controlled Protein Loading and Delivery. Adv Sci (Weinheim, Baden-Wurttemberg, Ger. 2023;10(34):e2304389. doi:10.1002/advs.202304389

17. Roefs MT, Gamauf J, Kroenigsberger B, et al. Rapid extracellular vesicle surface decoration with targeting moieties based on a fluorescein binding single chain variable fragment snorkel. J Control release Off J Control Release Soc. 2026;390:114558. doi:10.1016/j.jconrel.2025.114558

18. Zahradník J, Dey D, Marciano S, et al. A Protein-Engineered, Enhanced Yeast Display Platform for Rapid Evolution of Challenging Targets. ACS Synth Biol. 2021;10(12):3445–3460. doi:10.1021/acssynbio.1c00395

19. Götzke H, Kilisch M, Martínez-Carranza M, et al. The ALFA-tag is a highly versatile tool for nanobody-based bioscience applications. Nat Commun. 2019;10(1):4403. doi:10.1038/s41467-019-12301-7

20. Stadler C, Rexhepaj E, Singan VR, et al. Immunofluorescence and fluorescent-protein tagging show high correlation for protein localization in mammalian cells. Nat Methods. 2013;10(4):315–323. doi:10.1038/nmeth.2377

21. Hoffmann C, Gaietta G, Bünemann M, et al. A FlAsH-based FRET approach to determine G protein–coupled receptor activation in living cells. Nat Methods. 2005;2(3):171–176. doi:10.1038/nmeth742

22. Welsh JA, Goberdhan DCI, O’Driscoll L, et al. Minimal information for studies of extracellular vesicles (MISEV2023): From basic to advanced approaches. J Extracell vesicles. 2024;13(2):e12404. doi:10.1002/jev2.12404

23. Tian T, Zhang H-X, He C-P, et al. Surface functionalized exosomes as targeted drug delivery vehicles for cerebral ischemia therapy. Biomaterials. 2018;150:137–149. doi:10.1016/j.biomaterials.2017.10.012

24. d’Avanzo N, Torrieri G, Figueiredo P, et al. LinTT1 peptide-functionalized liposomes for targeted breast cancer therapy. Int J Pharm. 2021;597:120346. doi:10.1016/j.ijpharm.2021.120346

25. Gupta D, Wiklander OPB, Görgens A, et al. Amelioration of systemic inflammation via the display of two different decoy protein receptors on extracellular vesicles. Nat Biomed Eng. 2021;5(9):1084–1098. doi:10.1038/s41551-021-00792-z

26. Liang X, Niu Z, Galli V, et al. Extracellular vesicles engineered to bind albumin demonstrate extended circulation time and lymph node accumulation in mouse models. J Extracell vesicles. 2022;11(7):e12248. doi:10.1002/jev2.12248

27. Zheng W, Rädler J, Sork H, et al. Identification of scaffold proteins for improved endogenous engineering of extracellular vesicles. Nat Commun. 2023;14(1):4734. doi:10.1038/s41467-023-40453-0

28. Galisova A, Zahradnik J, Allouche-Arnon H, et al. Genetically Engineered MRI-Trackable Extracellular Vesicles as SARS-CoV-2 Mimetics for Mapping ACE2 Binding In Vivo. ACS Nano. 2022;16(8):12276–12289. doi:10.1021/acsnano.2c03119

29. Zhu Z, Zhai Y, Hao Y, et al. Specific anti-glioma targeted-delivery strategy of engineered small extracellular vesicles dual-functionalised by Angiopep-2 and TAT peptides. J Extracell vesicles. 2022;11(8):e12255. doi:10.1002/jev2.12255

30. Sun B, Lv Z, Li R, et al. Synergistic Dual-Targeting Bacterial Extracellular Vesicles Delivering Paclitaxel for Precision Glioblastoma Therapy. Nano Lett. 2025;25(43):15468–15477. doi:10.1021/acs.nanolett.5c03032

31. Yan L, Rosen N, Arteaga C. Targeted cancer therapies. Chin J Cancer. 2011;30(1):1–4. doi:10.5732/cjc.010.10553

32. Ingato D, Edson JA, Zakharian M, Kwon YJ. Cancer Cell-Derived, Drug-Loaded Nanovesicles Induced by Sulfhydryl-Blocking for Effective and Safe Cancer Therapy. ACS Nano. 2018;12(9):9568–9577. doi:10.1021/acsnano.8b05377

33. Debele TA, Mekuria SL, Tsai HC. Polysaccharide based nanogels in the drug delivery system: Application as the carrier of pharmaceutical agents. Mater Sci Eng C. 2016;68:964–981. doi:10.1016/j.msec.2016.05.121

34. Gálisová A, Jirátová M, Rabyk M, et al. Glycogen as an advantageous polymer carrier in cancer theranostics: Straightforward in vivo evidence. Sci Rep. 2020;10(1). doi:10.1038/s41598-020-67277-y

35. Wauters AC, Scheerstra JF, van Leent MMT, et al. Polymersomes with splenic avidity target red pulp myeloid cells for cancer immunotherapy. Nat Nanotechnol. 2024;19(11):1735–1744. doi:10.1038/s41565-024-01727-w

36. Peleg Y, Unger T. Application of the Restriction-Free (RF) cloning for multicomponents assembly. Methods Mol Biol. 2014;1116:73–87. doi:10.1007/978-1-62703-764-8_6

37. Zahradník J, Kolářová L, Peleg Y, et al. Flexible regions govern promiscuous binding of IL-24 to receptors IL-20R1 and IL-22R1. FEBS J. 2019;286(19):3858–3873. doi:10.1111/febs.14945

38. Shaner NC, Lambert GG, Chammas A, et al. A bright monomeric green fluorescent protein derived from Branchiostoma lanceolatum. Nat Methods. 2013;10(5):407–409. doi:10.1038/nmeth.2413

39. Shaner NC, Campbell RE, Steinbach PA, Giepmans BNG, Palmer AE, Tsien RY. Improved monomeric red, orange and yellow fluorescent proteins derived from Discosoma sp. red fluorescent protein. Nat Biotechnol. 2004;22(12):1567–1572. doi:10.1038/nbt1037

40. Pedersen ME, Østergaard J, Jensen H. Flow-Induced Dispersion Analysis (FIDA) for Protein Quantification and Characterization. Methods Mol Biol. 2019;1972:109–123. doi:10.1007/978-1-4939-9213-3_8

