## Supplementary Material for "Universal functionalization of extracellular vesicles with nanobody adapters"

Andrea Galisova[1,2*], Jiri Zahradnik[2], Ester Merunkova[1], Dominik Havlicek[1,3], Josef Uskoba[4], Ziv Porat[5], Daniel Jirak[1,3]

[1] Institute for Clinical and Experimental Medicine, Videnska 1958/9, Prague, Czech Republic
[2] The First Faculty of Medicine, Charles University, BIOCEV, Prumyslova 595, Vestec, Prague-region, Czech Republic
[3] Institute of Biophysics, The First Faculty of Medicine, Charles University, Salmovska 1, Prague, Czech Republic
[4] BioTech a.s., Kramolinska 955, Prague, Czech Republic
[5] Life Sciences Core Facilities, Weizmann Institute of Science, Herzl Street 234, Rehovot, Israel

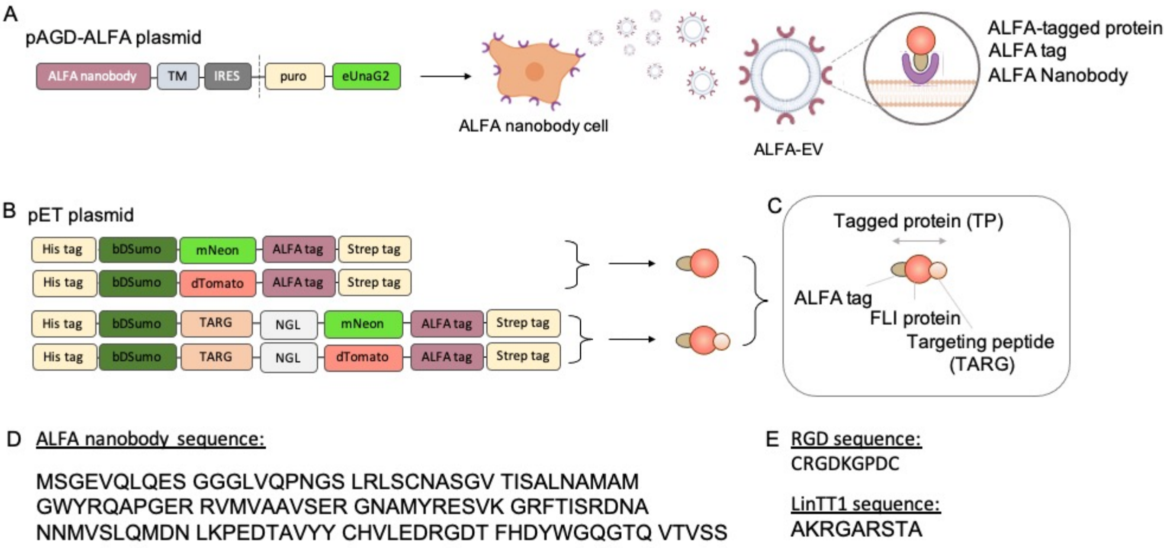


**Fig. S1 Preparation of genetic constructs for NaTaLi. (A)** A scheme of a plasmid for expression of ALFA nanobody on the cell membrane surface. TM stands for the PDGFR-based transmembrane domain. The resulting ALFA nanobody cell line serve as parental source for ALFA nanobody EVs that are isolated from these cells. **(B)** Schemes of pET plasmids used for preparation of tagged proteins with or without targeting peptides. **(C)** A scheme of a composition of tagged proteins (TP) with a targeting peptide (TARG). **(D)** Amino acid sequence of the ALFA nanobody and **(E)** tumor-targeting peptides.

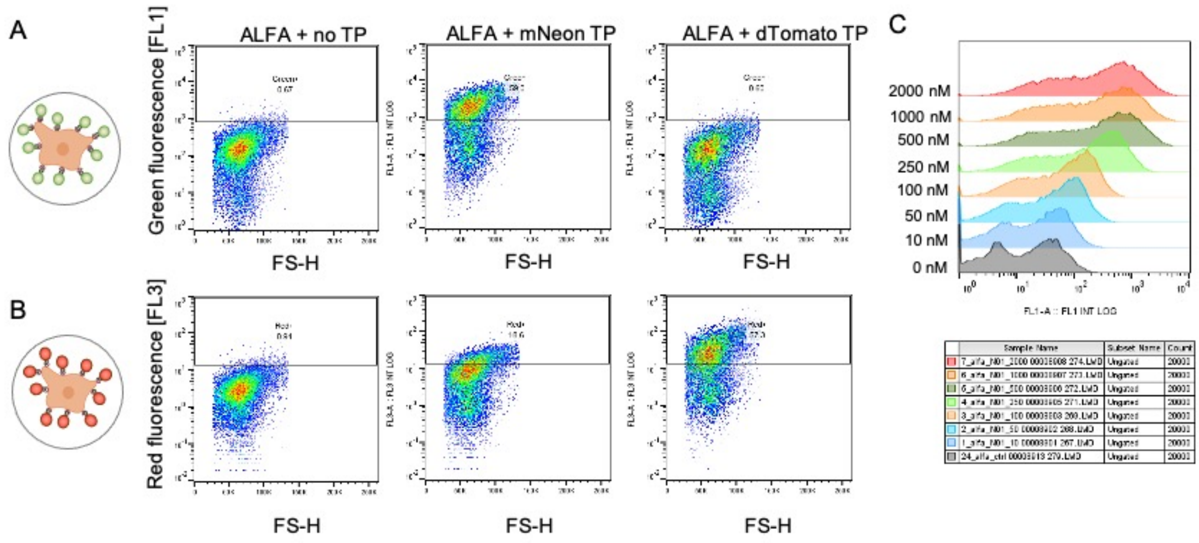


**Fig. S2 Characterization of parental ALFA-expressing cells.** Flow cytometry dotplots of cells functionalized with ALFA-tagged **(A)** mNeon or **(B)** dTomato. **(C)** Flow cytometry histograms of cells incubated with various concentration of ALFA-tagged mNeon protein used for quantification of the apparent equilibrium dissociation constant $K_D$.

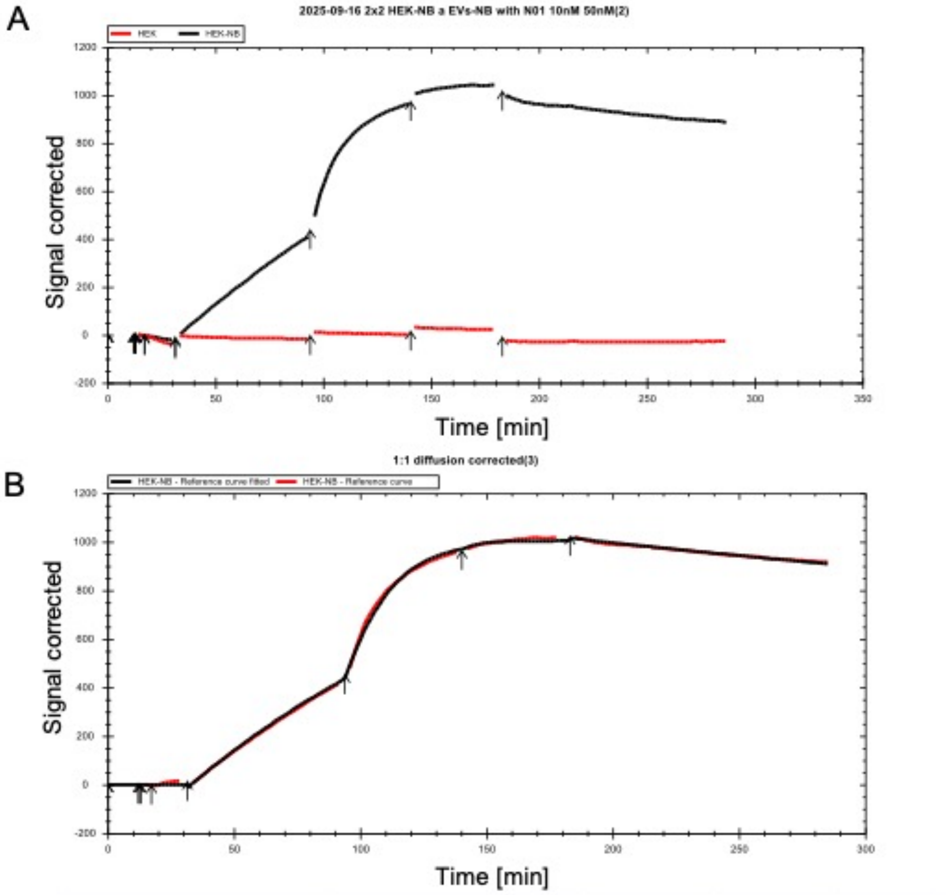


| Evaluation | Model | ka (1/(M*s)) | kd (1/s) | KD (M) |
|---|---|---|---|---|
| 1:1 diffusion corrected | Kinetics: 1:1 diffusion | 2,42E+04 | 1,89E-05 | 7,82E-10 |

**Fig. S3 Assessment of binding of the ALFA nanobody to tagged proteins. (A)** LigandTracer binding curve of ALFA-expressing cells incubated with mNeonTP (black line) and control protein (red line). **(B)** Fit of the binding curve with results.

**Purified proteins can be screened for binding to cancer biomarkers**

To ensure high purity, ALFA-tagged proteins were produced using an efficient and straightforward approach, involving overnight expression in E. coli BL21 (DE3) followed by gravity-flow tandem purification. This strategy employed His-tagged bdSUMO with on-column cleavage at the N-terminus and a Strep-tag at the C-terminus of the protein (Fig. S4A). Parental ALFA nanobody cells showed a distinct binding of RGD- and LinTT1-tagged proteins (Fig. S4C) proving proper binding of the ALFA tagged proteins to ALFA nanobodies *in vitro*. To screen the isolated tumor-binding tagged proteins for binding to tumor cell receptors, various cancer cell lines were incubated with tumor-targeted ALFA tagged proteins (RGD and LinTT1). All tumor cells (4T1, PC3, CHO) showed higher binding of tumor-targeted ALFA-tagged RGD and LinTT1 to cancer cells compared to controls (Fig. S4D).

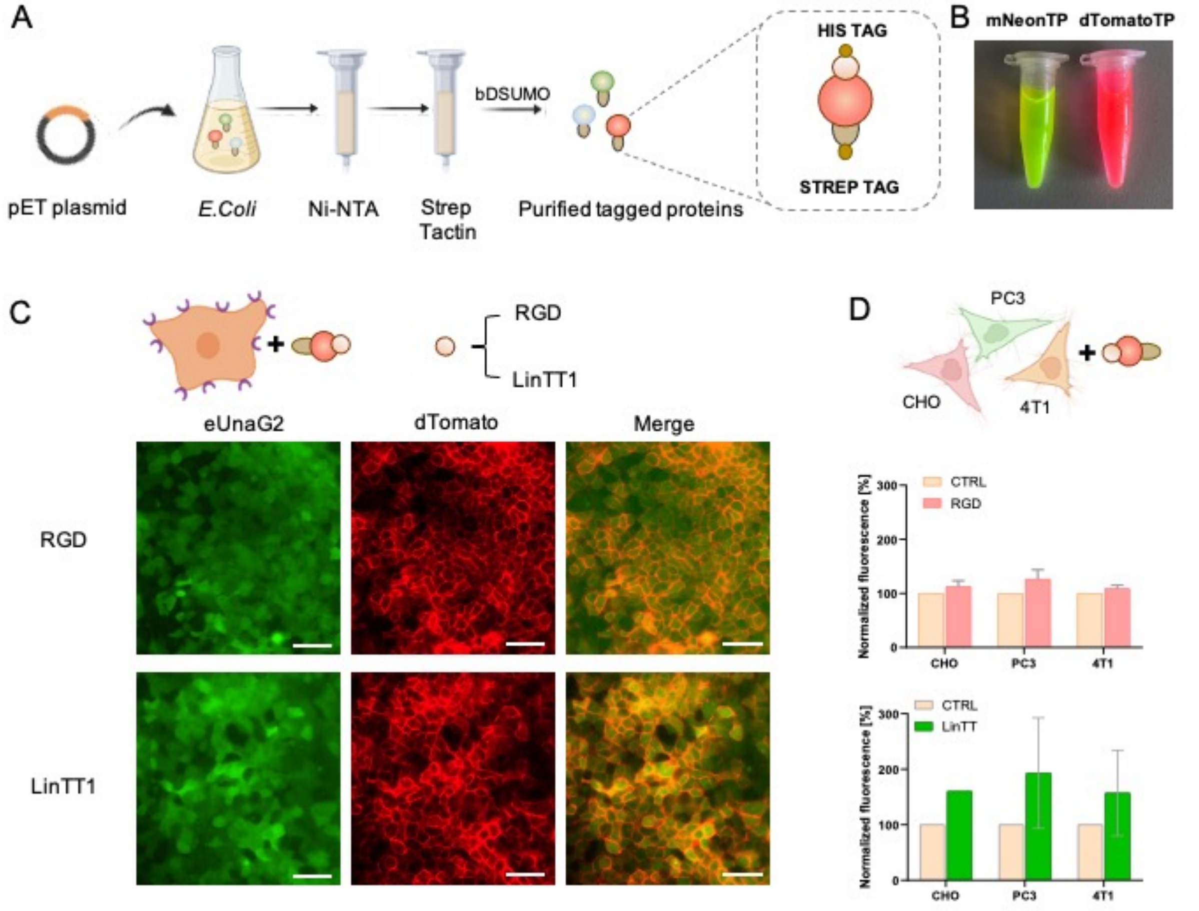


**Fig. S4 Production of proteins for ALFA-click-based conjugation to EVs and verification of their functionality. (A)** A scheme of production, isolation and purification of ALFA-tagged proteins from *E. coli*. **(B)** Photography of tubes with purified mNeon TP and dTomato TP. **(C)** Microscopy images of parental cells with bound tumor-targeting RGD and LinTT1 ALFA-tagged proteins. Scale bar is 200 µm. **(D)** Binding of tagged proteins RGD (up) and LinTT1 (down) to antigens on tumor cells measured by flow cytometry. Fluorescence was normalized to cells incubated without tagged proteins.

**Tandem purification of the tagged proteins**

Using the tandem purification protocol, the ALFA-tagged proteins of interest were more than 90% pure, exhibiting a single dominant band on SDS-PAGE (Fig. S5). In addition, tandem purification using tags positioned at the N- and C-termini ensures the integrity of the produced protein sequence and eliminates proteolytically degraded species. To reach the highest quality of proteins for functional in vitro and in vivo studies, RGD- and LinTT1-tagged proteins were additionally purified using size exclusion chromatography (Fig. S5F).

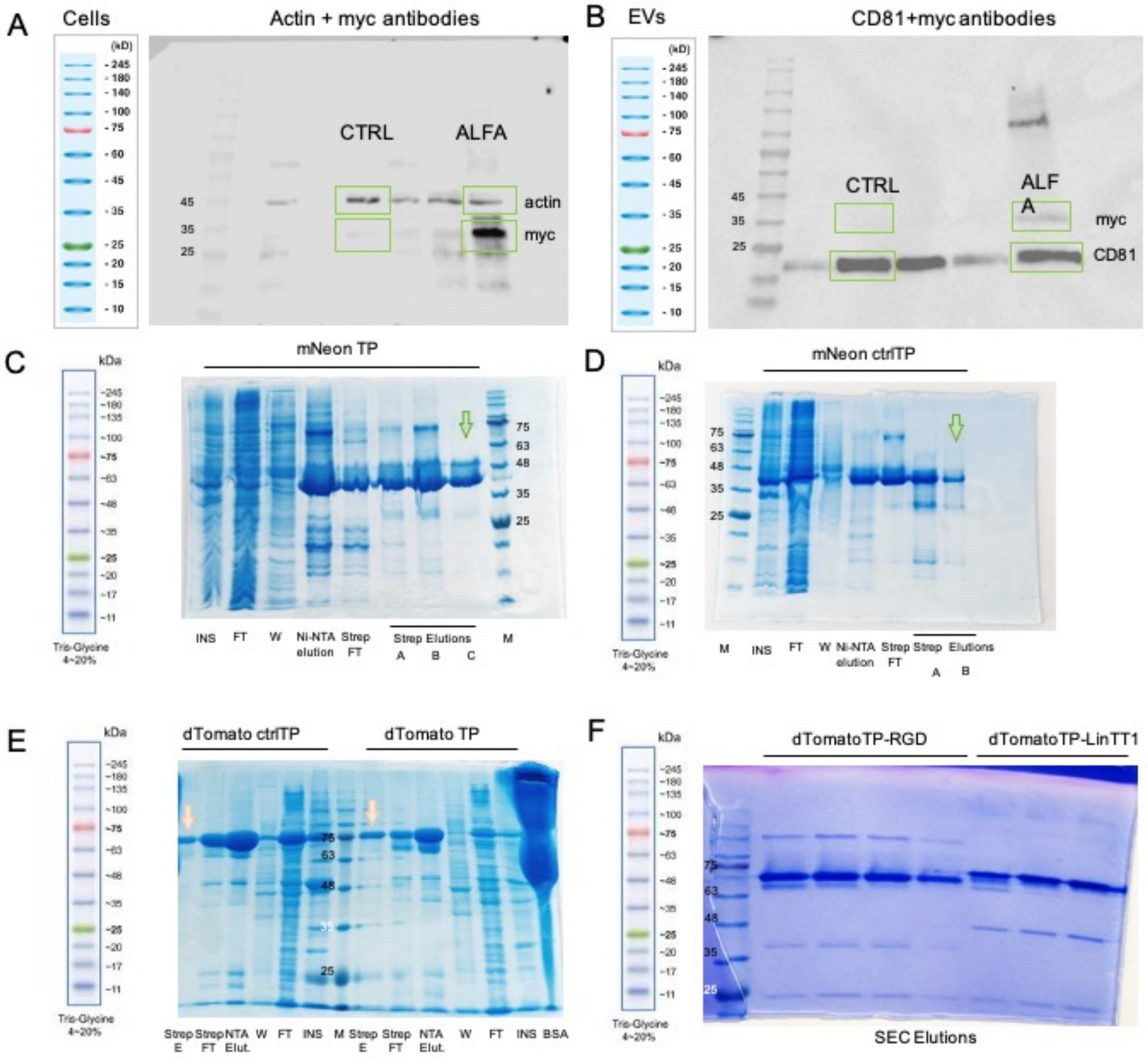


**Fig. S5 Characterization of purified proteins.** The full western blot membranes of (**A**) cell lysates and (**B**) EVs. Protein gels of **(C)** mNeonTP, **(D)** mNeon with ctrlTP, **(E)** dTomatoTP and dTomato with ctrpTP. **(F)** Protein gels of tagged proteins after size exclusion chromatography purification: dTomatoTP-RGD and dTomatoTP-LinTT1 protein. The arrows point to the final protein that was used for the follow-up experiments.

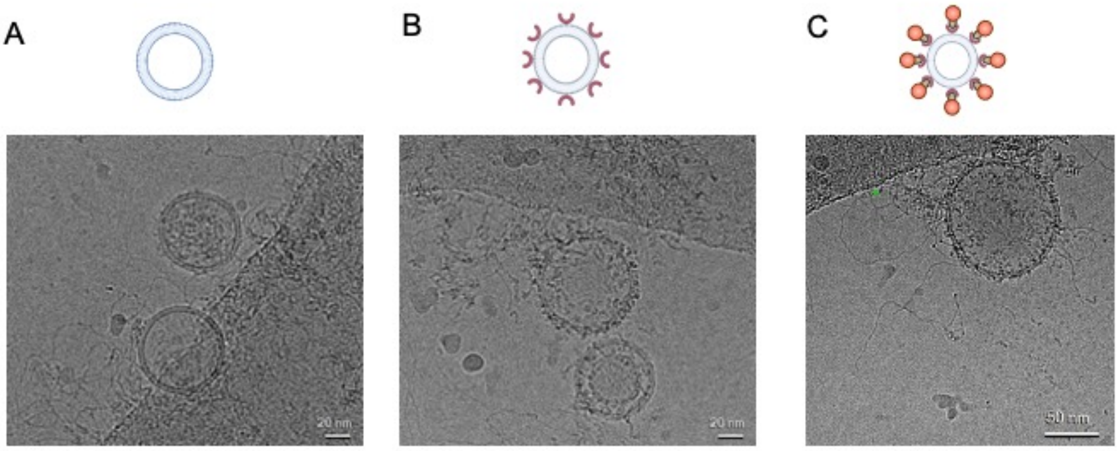


**Fig. S6 Cryo-EM images of EVs. (A)** Control EVs, **(B)** EVs with ALFA nanobodies and **(C)** functionalized EVs with nanobodies and clicked ALFA-tagged protein.

A

| Parameter Name | Unit | Value | Std. Dev. |
|---|---|---|---|
| Rh Indicator | nm | 3.4 | 0 |
| Rh Complex | nm | 82.12 | 5.536 |
| Kd | E+12 p_mL | 1.077E+9 | 2.667E+8 |

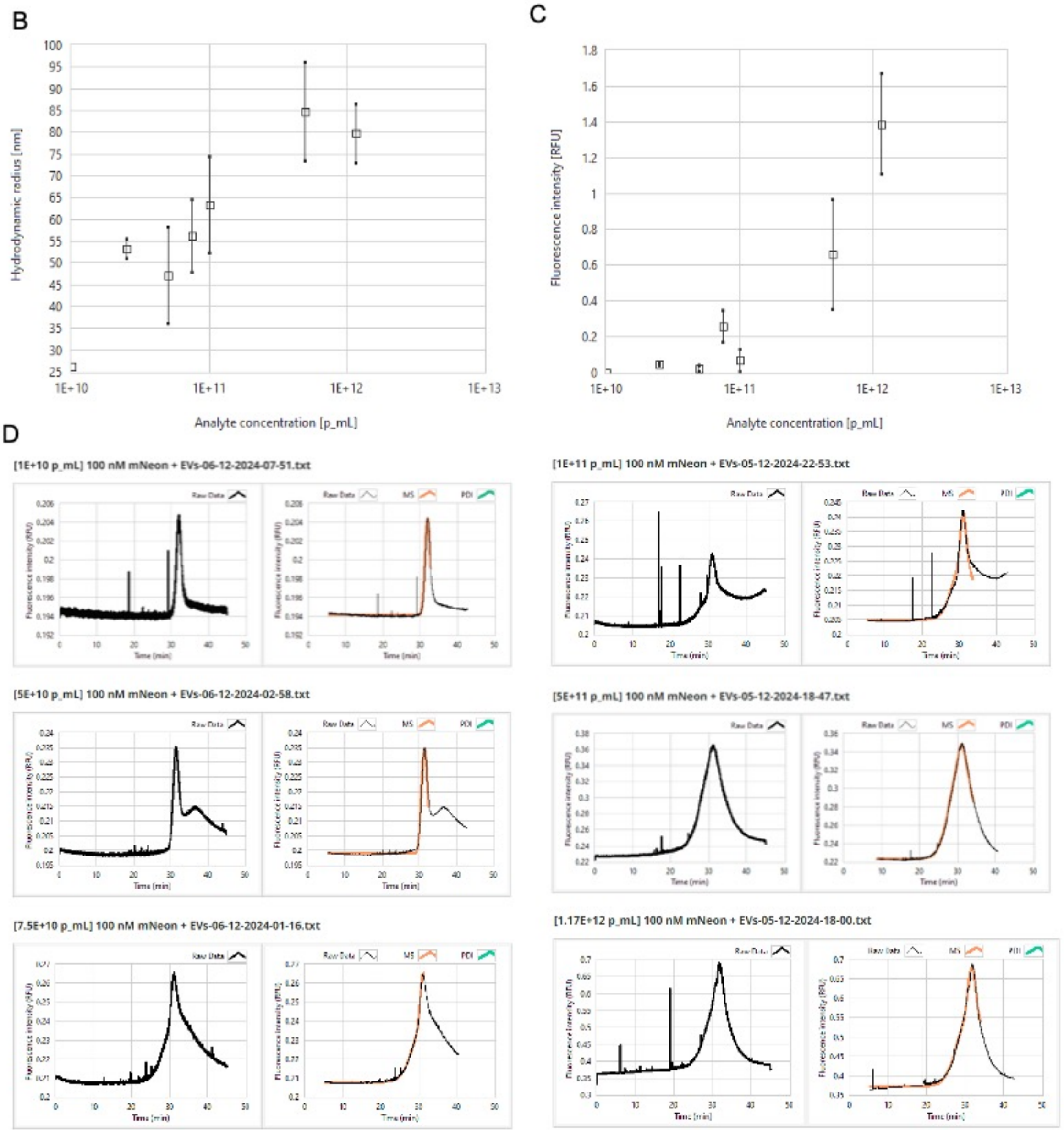


**Fig. S7 Flow Induced Dispersion Analysis (FIDA) of EVs. (A)** Hydrodynamic size of the ALFA-tagged mNeon protein (Rh indicator), ALFA-displaying EVs (Rh complex) and $K_D$ of the EVs-mNeon. **(B)** The change of fluorescence intensity in dependence to analyte concentration (EVs). **(C)** The change of hydrodynamic radius of EVs after binding with ALFA-tagged mNeon. **(D)** Individual fluorescence spectra for each EV concentration.

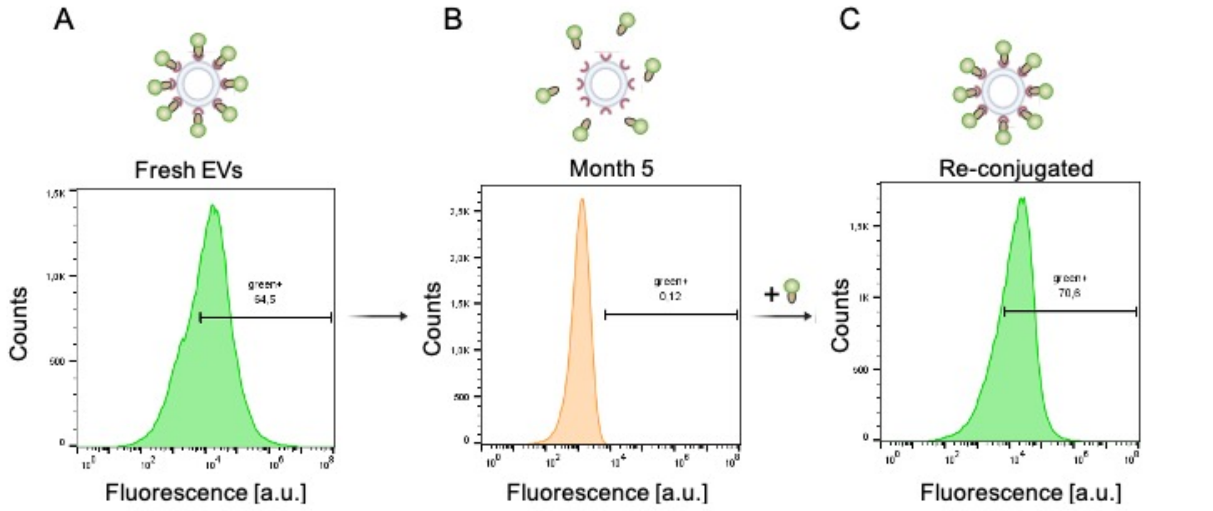


**Fig. S8 Stability of ALFA-EVs assessed by nanoflow cytometry.** Fluorescence signal of **(A)** ALFA-EVs with clicked mNeon protein immediately after isolation, **(B)** ALFA-EVs with mNeonTP 5 months after isolation functionalization and **(C)** freshly-reconjugated ALFA-EVs with mNeonTP (C).

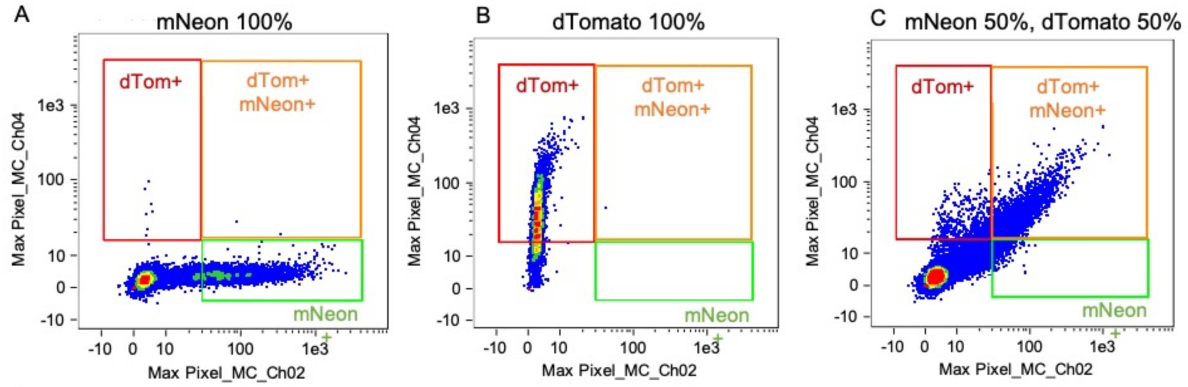


D

mNeonTP 100%

| Population | Count | %Gated | Objects/mL | Mean | Median | Std. Dev. |
|---|---|---|---|---|---|---|
| non-beads | 12444 | 100 | 14993519.07 | 46.74, 3.242 | 6.484, 3.225 | 104.46, 2.152 |
| mCherry + & non-beads | 7 | 0.06 | 8434.16 | 4.135, 48.32 | 4.014, 39.24 | 0.973, 28.95 |
| GFP + & non-beads | 4302 | 34.6 | 5183391.11 | 122.7, 4.127 | 74.86, 4.016 | 150.37, 1.418 |
| GFP+ mCherry+ & non-beads | 2 | 0.02 | 2409.76 | 205.58, 23.61 | 205.58, 23.61 | 168.98, 6.258 |

dTomatoTP 100%

| Population | Count | %Gated | Objects/mL | Mean | Median | Std. Dev. |
|---|---|---|---|---|---|---|
| non-beads | 7746 | 100 | 2964431.77 | 2.619, 43.5 | 2.528, 30.23 | 1.272, 50.63 |
| mCherry + & non-beads | 6112 | 78.9 | 2339092.04 | 2.769, 52.21 | 2.719, 37.6 | 1.174, 53.7 |
| GFP + & non-beads | 0 | 0 | 0 | NaN, NaN | NaN, NaN | NaN, NaN |
| GFP+ mCherry+ & non-beads | 1 | 0.01 | 382.7 | 41.21, 46.86 | 41.21, 46.86 | 0, 0 |

mNeonTP 50% dTomatoTP 50%

| Population | Count | %Gated | Objects/mL | Mean | Median | Std. Dev. |
|---|---|---|---|---|---|---|
| non-beads | 30667 | 100 | 10681269.08 | 11.18, 7.134 | 3.462, 3.183 | 30.84, 18.03 |
| mCherry + & non-beads | 626 | 2.04 | 218034.84 | 17.32, 27.09 | 18.35, 21.68 | 7.566, 14.8 |
| GFP + & non-beads | 492 | 1.6 | 171362.85 | 39.86, 12.72 | 36.91, 13.2 | 10.13, 2.573 |
| GFP+ mCherry+ & non-beads | 2116 | 6.9 | 736999.56 | 89.12, 48 | 65.34, 34.3 | 80.63, 50.57 |

**Fig. S9 Assessment of functionalization of EVs by ImageStream analysis.** Flow cytometry dotblots of **(A)** ALFA-EVs were functionalized with only mNeonTP, **(B)** only dTomatoTP and **(C)** at 50:50 ration of mNeonTP and dTomatoTP. **(D)** Analysis of the ImageStream signals of each EV formulation.

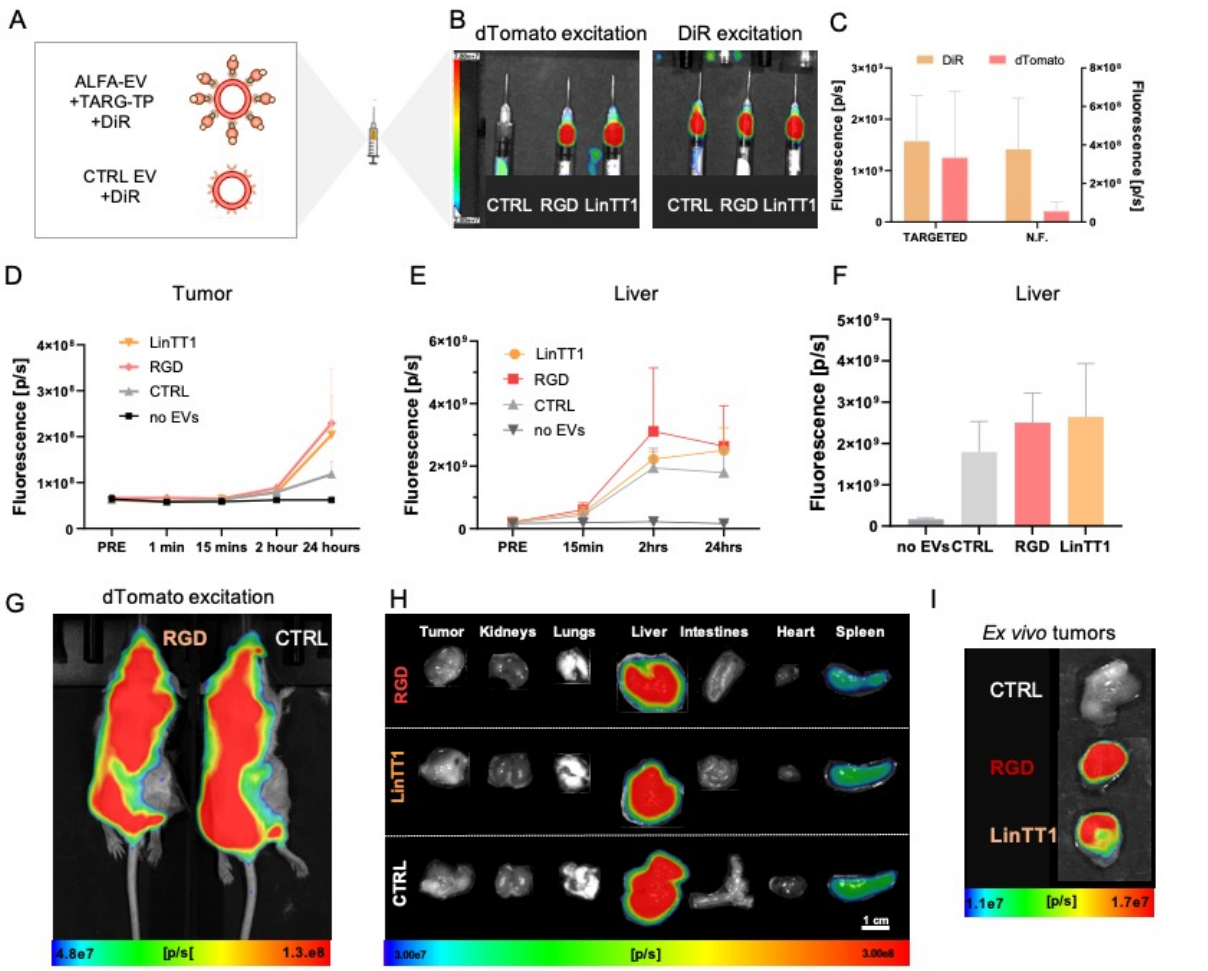


**Fig. S10 Tumor-targeted EVs. (A)** Functionalized ALFA-EVs with targeting peptide (EV-Nb-TARG-TP) and non-functionalized control EVs were labeled with a fluorescent dye DiR. **(B)** Fluorescence images of syringes filled with EVs for in vivo application after excitation at dTomato and DiR wavelengths. **(C)** Quantification of fluorescence signal from EVs in syringes. **(D)** *In vivo* changes of fluorescence signals over time in tumors after administration of EV formulations. **(E)** *In vivo* changes of fluorescence signals over time in the livers after administration of EV formulations. **(F)** Accumulation of intravenously administered EVs in the liver at the time point of 24 h after injection. **(G)** Fluorescence images of mice after administration of ALFA-EVs measured at dTomato excitation wavelengths. **(H)** Fluorescence images of tumors, kidneys, lungs, livers, intestines, hearts, spleen with the same color scale. **(I)** *Ex vivo* fluorescence images of harvested tumors.
